# Dual control of MAPK activities by AP2C1 and MKP1 MAPK phosphatases regulates defence responses in Arabidopsis

**DOI:** 10.1101/2021.06.03.446911

**Authors:** Zahra Ayatollahi, Vaiva Kazanaviciute, Volodymyr Shubchynskyy, Kotryna Kvederaviciute, Manfred Schwanninger, Wilfried Rozhon, Michael Stumpe, Felix Mauch, Sebastian Bartels, Roman Ulm, Salma Balazadeh, Bernd Mueller-Roeber, Irute Meskiene, Alois Schweighofer

## Abstract

Mitogen-activated protein kinase (MAPK) cascades transmit environmental signals and induce stress and defence responses in plants. These signalling cascades are negatively controlled by specific phosphatases of the type 2C Ser/Thr protein phosphatase (PP2C) and dual-specificity phosphatase (DSP) families that inactivate stress-induced MAPKs; however, the interplay between phosphatases of these different types has remained unknown. Our work reveals that different Arabidopsis MAPK phosphatases, the PP2C-type AP2C1 and the DSP-type MKP1, exhibit both specific and overlapping functions in plant stress responses. Each single mutant and *ap2c1 mkp1* double mutant displayed enhanced wound-induced activation of MAPKs MPK3, MPK4, and MPK6, as well as induction of a set of transcription factors. Moreover, *ap2c1 mkp1* double mutants show an autoimmune-like response, associated with elevated levels the stress hormones salicylic acid and ethylene, and of the phytoalexin camalexin. Interestingly, this phenotype is reduced in *ap2c1 mkp1 mpk6* triple mutants, suggesting that the autoimmune-like response is due to MPK6 misregulation. We conclude that the evolutionarily distant MAPK phosphatases AP2C1 and MKP1 contribute crucially to the tight control of MPK6 activity, ensuring appropriately balanced stress signalling and suppression of autoimmune-like responses during plant growth and development.

**Highlight:** Double MAPK phosphatase mutant plants *ap2c1 mkp1* exhibit constitutive, autoimmune-like stress responses, dependent on their substrate MAPK MPK6.

## Introduction

Reversible protein phosphorylation is one of the most commonly used mechanisms for the molecular transmission of stress signals and developmental cues. This mechanism is based on the opposing actions of protein kinases and protein phosphatases. Mitogen-activated protein kinases (MAPKs) are highly conserved major components of developmental and stress signalling cascades in eukaryotes. MAPKs are activated by upstream MAPK kinases via phosphorylation of Thr and Tyr within their activation loop. This activation eventually leads to the reprogramming of cellular activities, including the modulation of gene expression, to generate appropriate responses. The activation of MAPKs does not represent a simple on/off switch, as both the magnitude and duration of activation are crucial for determining the signalling outcome (Marshall, 1995). Prolonged or constant activation of a MAPK cascade can have detrimental effects as illustrated by the hypersensitive response (HR)-induced cell death in plants expressing a constitutively active MAPK kinase version (Liu *et al*., 2007; Ren *et al*., 2002). Thus, negative regulation and inactivation mechanisms are important for the correct cellular response. Specific protein phosphatases can dephosphorylate and thereby inactivate MAPKs. As dual phosphorylation of the Thr-X-Tyr motif in the activation loop is required for MAPK activation (Caunt and Keyse, 2013), dephosphorylation of either phospho-amino acid residue inactivates the MAPK and inhibits downstream signalling. Interestingly, this inactivation can be accomplished by evolutionarily distant protein phosphatases, including PP2C-type MAPK phosphatases (Fuchs *et al*., 2013; Schweighofer *et al*., 2004; Schweighofer *et al*., 2007) and PTP-type dual specificity (Tyr and Ser/Thr) phosphatases (DSPs) (Bartels *et al*., 2010; Jiang *et al*., 2018). However, their interplay is presently unknown.

*Arabidopsis thaliana* DSP-type mitogen-activated protein kinase phosphatase 1 (MKP1) interacts with the stress-responsive MAPKs MPK3, MPK4 and MPK6, and controls their activities (Anderson *et al*., 2011; Bartels *et al*., 2009; Ulm *et al*., 2002). The *mkp1* knockout mutant is hypersensitive to genotoxic stress, including UV-B radiation (Gonzalez Besteiro *et al*., 2011; Gonzalez Besteiro and Ulm, 2013; Ulm *et al*., 2002; Ulm *et al*., 2001), but is more resistant than wild type (WT) to the virulent bacterial pathogen *Pseudomonas syringae* pv. *tomato* (*Pst*) (Anderson *et al*., 2011; Anderson *et al*., 2014; Bartels *et al*., 2009). Specifically in the Arabidopsis Columbia accession, *mkp1* shows an autoimmune-like growth phenotype dependent on the disease resistance gene homologue *SUPPRESSOR OF npr1-1, CONSTITUTIVE 1* (*SNC1*) and is associated with enhanced MAPK activities (Bartels *et al*., 2009). The phospho-Tyr-specific PTP-type protein phosphatase PTP1 also interacts with MPK6 and MPK3 in transient assays. The lack of both MKP1 and PTP1 in the *mkp1 ptp1* double mutant leads to upregulation of MPK6-dependent plant defence responses and a further enhanced autoimmune-like phenotype (Bartels *et al*., 2009).

A group of PP2C-type phosphatases, including AP2C1, interacts with MAPKs and controls their activities (Schweighofer *et al*., 2007; Umbrasaite *et al*., 2010). AP2C1 is induced by wounding and biotic stress, and functions as a negative regulator of MPK3, MPK4 and MPK6 controlling levels of wound-induced jasmonate and ethylene (ET) as well as plant immunity (Galletti *et al*., 2011; Schweighofer *et al*., 2007; Shubchynskyy *et al*., 2017; Sidonskaya *et al*., 2016). *ap2c1* plants do not display obvious developmental phenotypes under standard growth conditions (Schweighofer *et al*., 2007), implying specific AP2C1 function under stress conditions and contribution of other, presently unknown, MAPK phosphatases for MAPK control in the absence of AP2C1.

Activation of transcription factors (TFs) and changes of gene expression are part of the cellular response to a perceived signal in order to reprogram cellular processes (Rauf *et al*., 2013). A number of TFs, including WRKY and AP2-domain/ethylene-responsive factor (AP2/ERF) family members, have been suggested or demonstrated to act downstream of MAPKs in plants (Asai *et al*., 2002; Bethke *et al*., 2009; Guan *et al*., 2014; Kim and Zhang, 2004; Li *et al*., 2012; Mao *et al*., 2011; Meng and Zhang, 2013; Menke *et al*., 2005; Nakano *et al*., 2006; Popescu *et al*., 2009). Subsequently, these proteins may constitute an important link between pathogen-or wound-induced MAPK signalling and downstream transcriptional reprogramming.

Considering the broad spectrum of signals transmitted by the same MAPKs (Meng and Zhang, 2013; Rodriguez *et al*., 2010), such as MPK6, it is puzzling how the specificity of the responses for perceived stimuli is generated. The phylogenetic diversity and distinct enzymatic mechanisms of protein phosphatases that are able to inactivate MAPKs support the idea of a contribution of MAPK phosphatases to the versatility and specificity of MAPK networks. Here, we investigate the roles of the phylogenetically distant Arabidopsis MAPK phosphatases AP2C1 and MKP1 and, in particular, their functional redundancies. We show that AP2C1 and MKP1 together repress plant autoimmune-like responses, including salicylic acid (SA) and ET accumulation, and early senescence. These observations in the *ap2c1 mkp1* mutant are underlined by the misexpression of specific transcription factors, including members of the *WRKY, AP2/ERF*, and Arabidopsis NAM, ATAF, and CUC (*ANAC*) families whose expression is – at least partially – mediated by MPK6.

## Materials and methods

### Plant lines, genetic crosses and growth conditions

All plant lines used in this study were in the *Arabidopsis thaliana* accession Columbia (Col-0), with *mkp1* being an introgression line from a Wassilewskija background (Bartels *et al*., 2009). The T-DNA insertion line *ap2c1* (SALK_065126; (Schweighofer *et al*., 2007)) was crossed with the T-DNA insertion lines *ptp1* (SALK_118658) and *mkp1*, respectively, to generate the *ap2c1 ptp1* and *ap2c1 mkp1* double mutants*. mpk6-2* (SALK_073907) was used for genetic crosses generating *ap2c1 mkp1 mpk6.* The T-DNA insertion lines *ap2c2* (GABI-Kat_316F11) and *ap2c3* (SALK_109986) (Umbrasaite *et al*., 2010) were crossed with *mkp1* to generate *ap2c2 mkp1* and *ap2c3 mkp1* double mutants. Combinatorial mutants were identified in the F2 generation and also confirmed in subsequent generations by PCR genotyping using T-DNA- and gene-specific primers (Bartels *et al*., 2009; Schweighofer *et al*., 2007; Umbrasaite *et al*., 2010). For protein and RNA extraction, as well as for ET and SA measurements, plants were grown on soil for five to seven weeks in a phytotron chamber under short-day conditions (8 h light, 22°C/16 h dark, 20°C cycle).

For experiments at the seedling stage, seeds were surface sterilized and spread on plates containing half-strength MS (Murashige and Skoog) medium (Duchefa), pH 5.7, 1% (w/v) sucrose and 0.7% plant agar (w/v; Duchefa). Seedlings were grown in long-day conditions (16 h light/8 h dark) at 22°C. If indicated, *ap2c1 mkp1* plants were kept at 26°C during day and 22°C at night with 95% humidity and short-day conditions (8 h light/16 h dark).

### *Ex vivo* kinase activity assay and MAPK immunoblotting

Plant protein extraction and the *ex vivo* kinase assay were performed as described (Schweighofer *et al*., 2007; Schweighofer *et al*., 2009) using polyclonal antibodies for immunoprecipitation and myelin basic protein (MBP) as *in vitro* substrate of immunoprecipitated MAPKs. MAPK protein amounts were visualised with Sigma antibodies Anti-AtMPK3 (M8318), Anti-MPK4 (A6979) and Anti-AtMPK6 (A7104).

### RNA extraction and quantitative reverse-transcription PCR (RT-qPCR)

Total RNA from leaves was isolated with the RNeasy Plant Mini Kit (Qiagen) and treated with TURBO DNA-free DNaseI (Ambion) according to the manufacturers’ instructions. RNA integrity was checked on 1% (w/v) agarose gels and the concentration measured before and after DNAse I digestion. The absence of genomic DNA was verified by PCR using primers targeting an intron of the control gene *At5g65080*. cDNA synthesis was performed using First Strand cDNA Synthesis Kit (Thermo Scientific). The efficiency of cDNA synthesis was estimated by RT-qPCR analysis using a primer pair amplifying the 3’ part of the control gene encoding GAPDH and a primer pair amplifying the 5’ part of the same gene. RT-qPCR reactions were performed as described previously (Balazadeh *et al*., 2008). *ACTIN2* was selected as a reference gene for which four replicates were measured in each PCR run, and their average cycle threshold (CT) was used for relative expression analyses. TF expression data were normalized by subtracting the mean *ACTIN2* gene CT value from the CT value (ΔCT) of each gene of interest. The expression value in the comparison between different genotypes was calculated using the expression 2^-ΔΔCT^, where ΔΔCT represents ΔCT mutant of interest minus ΔCT control (wild type, WT). For TF expression profiling, an advanced version of an expression profiling platform (Balazadeh *et al*., 2008) that was originally described by (Czechowski *et al*., 2004) was used, covering 1,880 Arabidopsis TF genes. Statistical analysis was performed with the JASP software (https://jasp-stats.org; version 0.14.1).

### ET measurements, quantification of total SA and camalexin

ET measurements were performed by gas chromatography (Hewlett Packard 5890 Series II) with an Al_2_O_3_ column (Agilent Technologies). Whole rosettes of 4-week-old plants grown in long-day conditions were taken, leaves wounded, transferred into 20-mL vials containing 4 mL half-strength MS medium with 0.8% (w/v) plant agar, in order to reduce the volume of the head space, and air-tightly sealed. After 24 h, 100 µL of the gas phase were taken from the vials and analysed by gas chromatography–flame ionization detection (GC-FID). ET production was calculated per hour and milligram of fresh tissue.

Total SA was quantified as described previously (Rozhon *et al*., 2005) except that 20 µM EDTA was added to the HPLC eluent. Camalexin levels were determined as described previously (Shubchynskyy *et al*., 2017).

### AGI codes

The AGI codes of the genes analysed in this report are:

At1g01010, At1g02220, At1g02250, At1g02340, At1g04240, At1g04370,

At1g07160, At1g08320, At1g13300, At1g18570, At1g18860, At1g18860,

At1g19210, At1g29860, At1g36060, At1g52830, At1g62300, At1g66560,

At1g66560, At1g67100, At1g69600, At1g71520, At1g71860, At1g74080,

At1g75250, At1g80840, At2g01200, At2g30020, At2g33710, At2g37430,

At2g38250, At2g38340, At2g38470, At2g40180, At2g40470, At2g40740,

At2g40740, At2g43000, At2g43140, At2g43790, At2g47520, At3g01970,

At3g04070, At3g13840, At3g15320, At3g15500, At3g21330, At3g23230,

At3g23240, At3g23250, At3g26790, At3g26830, At3g44350, At3g45640,

At3g46080, At3g46090, At3g50510, At3g53600, At3g55270, At4g01370,

At4g01720, At4g01720, At4g18170, At4g18870, At4g23810, At4g32280,

At4g36990, At5g01900, At5g04390, At5g13080, At5g13330, At5g15160,

At5g19520, At5g20240, At5g22570, At5g23000, At5g24110, At5g27810,

At5g39860, At5g43290, At5g43650, At5g44260, At5g45890, At5g46350,

At5g51790, At5g56840, At5g56960, At5g59820, At5g64750, At5g64810,

At5g67450.

## Results

### Double *ap2c1 mkp1* mutant plants show growth and development defects which are at least partially mediated by MPK6

To investigate the specific and/or overlapping roles of the MAPK phosphatases AP2C1 and MKP1, we took advantage of the Arabidopsis T-DNA insertion knock-out mutants *ap2c1* and *mkp1*, respectively (Bartels *et al*., 2009; Schweighofer *et al*., 2007). Phenotypically, *ap2c1* and *mkp1* mutant plants did not show any difference compared to WT when grown for up to five weeks under short-day conditions (Figure 1A, 1C, 1E). However, long-day *mkp1* plants demonstrated altered morphology, such as aberrant leaf development and early senescence, which appeared approximately three weeks after germination (Supplementary Figure 1A), as described previously (Bartels *et al*., 2009).

**Figure 1.**
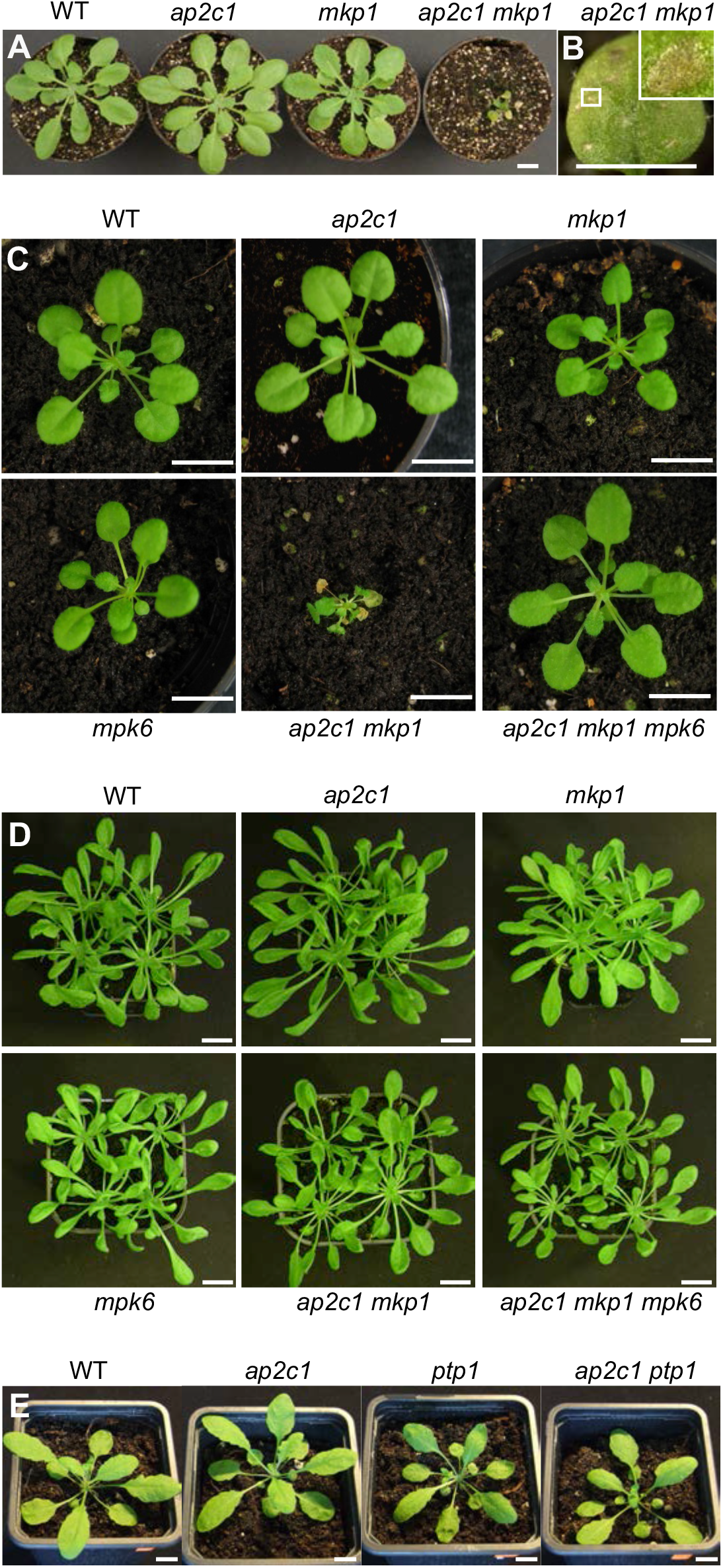
Loss of both AP2C1 and MKP1 causes developmental defects and precocious cell death, mediated by MPK6. **A.** Phenotypes of WT, *ap2c1*, *mkp1* and *ap2c1 mkp1* plants grown for 8 weeks in short-day conditions. Scale bars = 1 cm. **B.** Formation of macroscopic lesions in leaves of *ap2c1 mkp1* plant shown in Figure 1A. Scale bars = 0.5 cm. **C.** Phenotypes of five-week-old WT, *ap2c1*, *mkp1*, *mpk6*, *ap2c1 mkp1* and *ap2c1 mkp1 mpk6* plants grown in standard short-day conditions. Scale bars = 1 cm. **D.** Phenotypes of five-week-old WT, *ap2c1*, *mkp1*, *mpk6*, *ap2c1 mkp1* and *ap2c1 mkp1 mpk6* plants grown in conditions with increased humidity and elevated temperature. Scale bars = 1 cm. **E.** Phenotypes of WT, *ap2c1*, *ptp1* and *ap2c1 ptp1* plants grown for 6 weeks in short-day conditions. Scale bars = 1 cm.

To further analyse AP2C1 and MKP1 functions in plants, we generated a double mutant by genetic crossing. *ap2c1 mkp1* plants showed phenotypic differences compared to WT and single mutants. These appeared two weeks after germination under standard growth conditions in soil; first their sizes started differing, and during further growth *ap2c1 mkp1* plants revealed more pronounced multiple defects, including severe dwarfism and aberrant leaf development (Figures 1A, 1B, 1C). Four weeks after germination, phenotypic abnormalities became even more evident as early senescence, spontaneous macroscopic lesions and abnormal leaf morphology. Interestingly, these developmental defects were suppressed when plants were grown under conditions of elevated humidity and increased temperature (Figure 1D), indicating a dependency on environmental cues. However, during flowering, misshaped inflorescences and strongly reduced fertility were always observed (Supplementary Figure 2D, 2E). These phenotypes were specific for *ap2c1 mkp1* plants, as crossing *ap2c1* with *ptp1* did not lead to phenotypic alterations (Figure 1E) compared with *mkp1 ptp1* (Bartels *et al*., 2009). Crossing *mkp1* with either of two other clade B AP2C mutants, *ap2c2* or *ap2c3* (Umbrasaite *et al*., 2010), led to mild phenotypes compared with the strong defects of *ap2c1 mkp1* plants (Supplementary Figure 1B, C).

Since MPK6 is a documented target of action by AP2C1 and MKP1 (Schweighofer *et al*., 2007; Ulm *et al*., 2002) we set to address the impact of MPK6 on phenotypic aberrations detected in *ap2c1 mkp1* plants. To this goal, triple mutant plants *ap2c1 mkp1 mpk6* were created and their phenotype was compared with that of *ap2c1 mkp1*. The phenotype of *ap2c1 mkp1 mpk6* plants was more similar to WT than to *ap2c1 mkp1*. The loss of MPK6 suppressed most phenotypic defects observed in *ap2c1 mkp1*, such as extreme dwarfism, aberrant leaf shapes, premature leaf senescence and impaired fertility (Figure 1C). However, at later developmental stages, *ap2c1 mkp1 mpk6* plants appeared overall smaller than WT and displayed senescence in the older leaves (Supplementary Figures 1C, 2G).

Overall, these results suggest that AP2C1 and MKP1 protein phosphatases act partially redundantly and that the presence of at least either gene is necessary for normal plant development. The phenotypes observed in *ap2c1 mkp1* plants are predominantly MPK6-dependent.

### Dual control of wound-induced MAPK activities by AP2C1 and MKP1

Our previous work has revealed the involvement of AP2C1 in the regulation of MAPK activities induced by pathogen-associated molecular patterns (PAMP), wounding, and nematodes (Schweighofer *et al*., 2007; Shubchynskyy *et al*., 2017; Sidonskaya *et al*., 2016). To check for a potential overlapping role by MKP1, we firstly analysed MPK3, MPK4 and MPK6 activities after wounding the leaves of WT and of the single mutant plants *ap2c1* and *mkp1*. Kinase activities were assayed after immunoprecipitation from total protein extracts using specific antibodies. In agreement with our previous findings (Schweighofer *et al*., 2007), *ap2c1* plants showed higher and sustained wound-induced activities of MPK3, MPK4 and MPK6 compared to WT (Figure 2). Interestingly, MPK4 activity was more intense and sustained in *ap2c1* compared to the *mkp1* plants, indicating a specific role of AP2C1 in the regulation of MPK4 during wounding. MPK3 activity in *ap2c1* plants was slightly enhanced and more sustained in comparison to WT and *mkp1*. In *mkp1* plants, however, the MPK6 peak activity was shifted to an earlier time point compared to WT. In *ap2c1 mkp1* plants we detected strongly and moderately enhanced basal activity of MPK4 and MPK6, respectively, whereas basal activity of MPK3 was not affected in comparison to WT or single mutant lines. The stronger and more sustained wound-induced activation of MAPKs observed in single-mutant plants was additionally enhanced in the double mutant *ap2c1 mkp1* (Figure 2). The MPK3, MPK4, and MPK6 protein levels were comparable in both mutants and WT (Figure 2).

**Figure 2.**
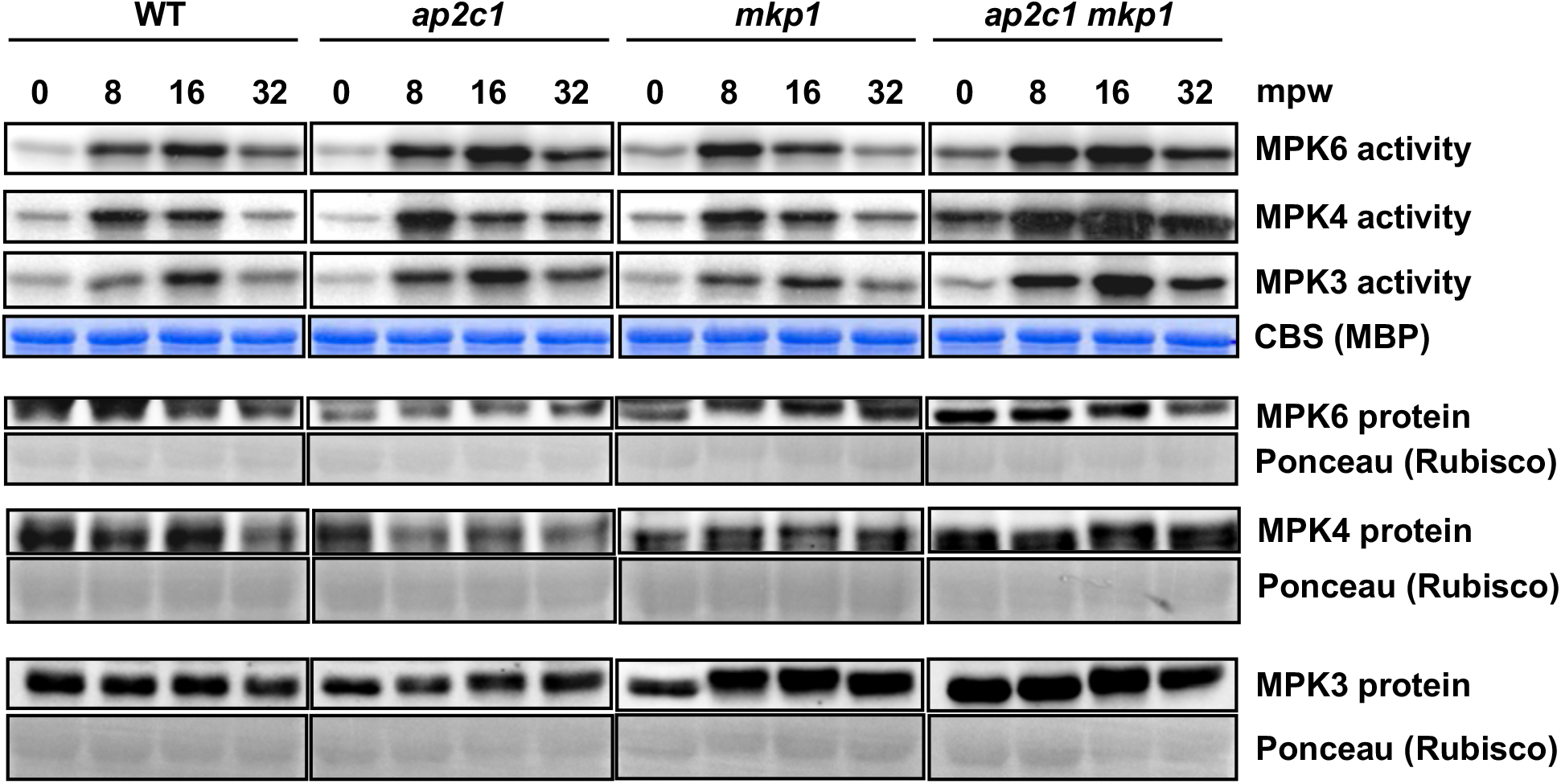
AP2C1 and MKP1 control wound-induced MAPK activities. Analysis of wound-induced MPK6, MPK4 and MPK3 kinase activities and protein amounts of leaves from six-week-old WT, *ap2c1, mkp1,* and *ap2c1 mkp1* plants grown in short-day conditions. MAPK activities were determined after immunoprecipitation by phosphorylation of myelin basic protein detected by autoradiography. Loading is demonstrated by Coomassie Blue staining (CBS); representative lanes are shown. MAPK protein amounts before and after wounding are demonstrated by immunoblotting of MPK3, MPK4, and MPK6 from total protein extract using specific antibodies. Loading is demonstrated by Ponceau S staining (Rubisco protein). mpw: minutes post wounding.

*MKP1* has been reported to be constitutively expressed (Ulm *et al*., 2002), whereas *AP2C1* is transcriptionally responsive to stress (Schweighofer *et al*., 2007; Sidonskaya *et al*., 2016). We tested whether reciprocal compensational expression may occur in long-day conditions and thus analysed *AP2C1* and *MKP1* mRNA levels in *mkp1* and *ap2c1* mutants, respectively. RT-qPCR analyses showed only very slightly enhanced expression of *MKP1* in *ap2c1* plants, whereas the expression of *AP2C1* was approximately 160% in *mkp1* plants compared to WT (Figure 3), suggesting a compensatory transcriptional activation of *AP2C1* in the absence of MKP1.

**Figure 3.**
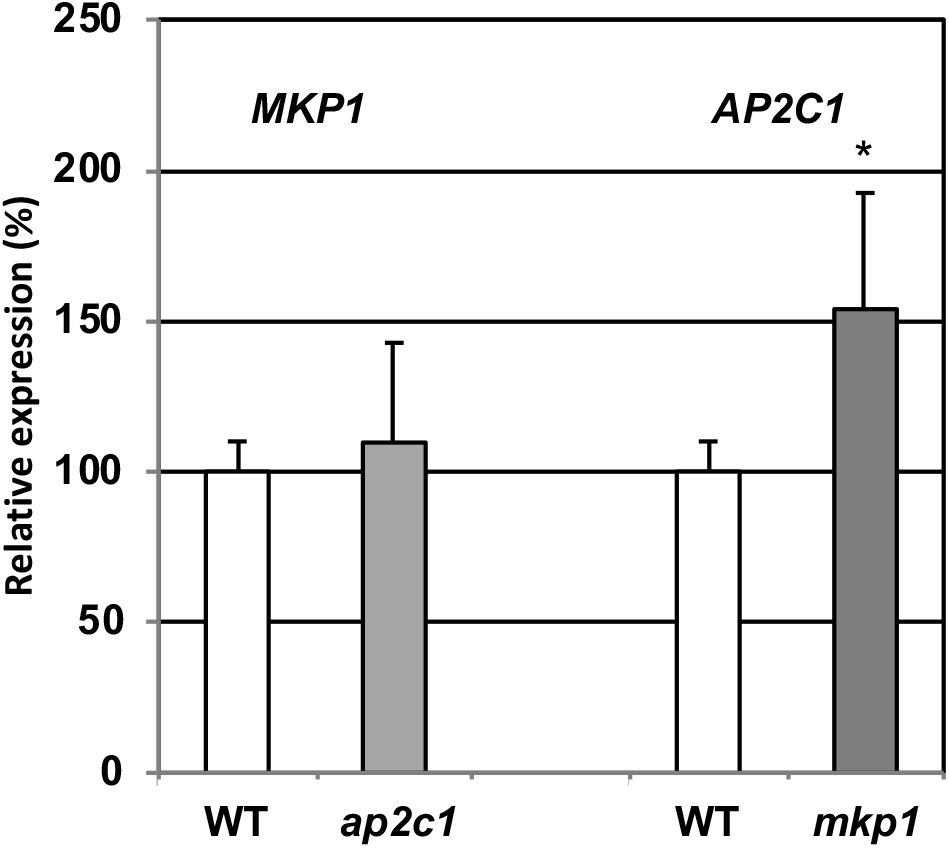
Detection of *MKP1* and *AP2C1* expression levels in *ap2c1* and *mkp1* mutants. Levels of *MKP1* and *AP2C1* transcripts were quantified by RT-qPCR in 14-day-old seedlings grown in long-day conditions, and compared with levels in WT plants. Transcript amounts of *MKP1* and *AP2C1* in WT were taken as 100%. The relative transcript amounts were normalized to the reference gene, *ACTIN2*. Results are the mean of two biological and two technical replicates for each experiment, *p < 0.05, Student’s *t*-test.

Our results suggest cooperative action and partial redundancy in the regulation of MAPKs by these two evolutionary distant and unrelated MAPK phosphatases.

### AP2C1 and MKP1 play partially redundant roles in the control of wound-induced ET synthesis

Enhanced ET production is an early response of plants subjected to biotic/abiotic stresses (Ju and Chang, 2012; Wang *et al*., 2002). We have previously shown that ectopic expression of *AP2C1* suppresses MPK6 activation and wound-induced ET production in plant leaves (Schweighofer *et al*., 2007). Since both AP2C1 and MKP1 control MPK6 activity, a major determinant in the regulation of ET biosynthesis (Li *et al*., 2012; Liu and Zhang, 2004), we analysed wound-induced ET amounts in leaves of WT, *ap2c1*, *mkp1*, *ap2c1 mkp1* and *ap2c1 mkp1 mpk6* plants. As reported earlier (Schweighofer *et al*., 2007), wound-induced ET amounts were similar in *ap2c1* and WT (Figure 4A). However, significantly higher levels of ET accumulated in wounded *mkp1* plants and even more so in the *ap2c1 mkp1* double mutant (Figure 4A). Our data suggest a primary role of MKP1 in the control of wound-triggered ET production and that although disruption of AP2C1 alone is not sufficient to alter ET production upon wounding, it contributes significantly to the regulation of ET amounts in the absence of MKP1. Interestingly and in agreement with the overall milder phenotype, wound-induced ET accumulation in *ap2c1 mkp1 mpk6* plants was similar to levels detected in WT (Figure 4A).

**Figure 4.**
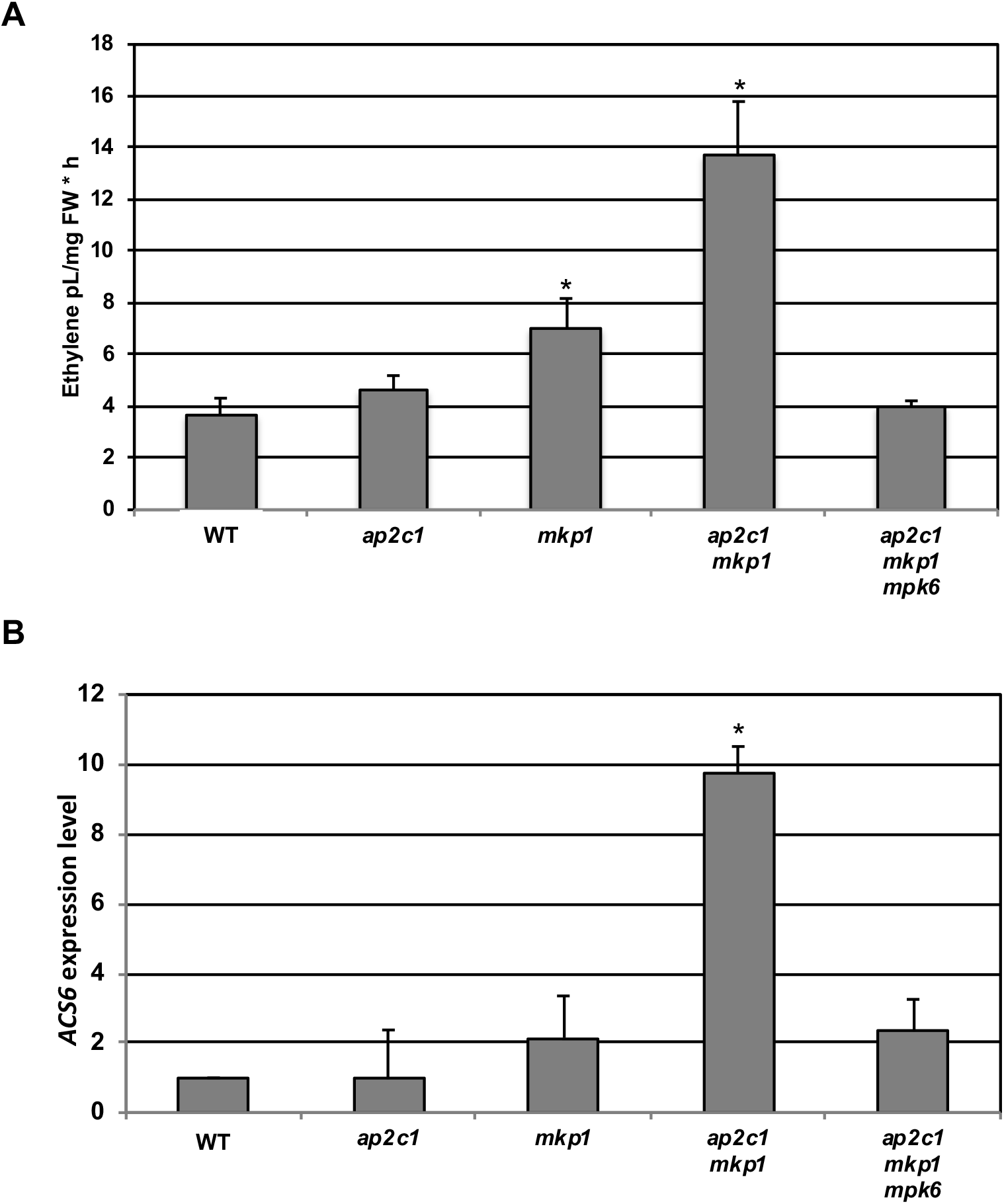
ap2c1 *mkp1* plants have elevated *ASC6* expression and produce more ET upon wounding than WT, mainly mediated by MPK6. **A.** ET levels produced by four-week-old WT, *ap2c1*, *mkp1, ap2c1 mkp1*, and *ap2c1 mkp1 mpk6* plants grown in standard long-day conditions. Values are expressed as ET amounts produced per mg plant fresh weight (FW) per hour. Bars represent mean values of three biological replicates ± SD, *p < 0.05 Student’s *t*-test. **B.** RT-qPCR analysis of *ASC6* expression in leaves of six-week-old WT, *ap2c1*, *mkp1, ap2c1 mkp1*, and *ap2c1 mkp1 mpk6* plants grown in short-day conditions, where expression levels in WT are set to 1. Bars represent mean values of three biological replicates ± SD, *p < 0.05 Student’s *t*-test.

The transcriptional regulation of 1-aminocyclopropane-1-carboxylic synthase (ACS) enzymes contributes to control ET production (Li *et al*., 2012). Therefore, we quantified the transcripts of *ACS6,* the expression of which is significantly induced after pathogen attack (Li *et al*., 2012) and wounding (Li *et al*., 2018). Compared to WT, no changes in *ACS6* transcript levels were detected in *ap2c1*, slightly higher levels in *mkp1*, and a nine-fold increase in *ap2c1 mkp1* which was reduced to WT levels in *ap2c1 mkp1 mpk6* plants (Figure 4B). Thus, our data show that *ACS6* is more expressed in *ap2c1 mkp1* plants, which likely contributes to the elevated amounts of ET upon wounding, and that both effects are mediated by MPK6.

Taken together, the wound-induced MAPK activities, expression patterns and effects on ET production suggest that AP2C1 and MKP1 have both distinct as well as overlapping functions in wounded leaves.

### TF gene expression is de-regulated in *ap2c1 mkp1* plants

To investigate if and how AP2C1 and MKP1 influence the regulation of gene expression under standard growth conditions, we used a RT-qPCR platform for high-throughput expression profiling of 1,880 Arabidopsis TF-encoding genes (Balazadeh *et al*., 2008). We selected genes showing an at least three-fold mean difference of expression levels in *ap2c1*, *mkp1* or *ap2c1 mkp1* plants when compared to WT. We identified three genes encoding TFs that were deregulated in *ap2c1*, but not in *mkp1* (Supplementary Table I), while 25 genes were deregulated in *mkp1*, but not in *ap2c1* (Supplementary Table II), and four genes concomitantly regulated by AP2C1 and MKP1 (Supplementary Table III). Figure 5 shows the number of genes whose expression levels were changed in *ap2c1*, *mkp1* or *ap2c1 mkp1* plants, compared to the WT. The TF genes dysregulated in the double mutant, and their expression values relative to the WT, are represented in Supplementary Table IV. The deregulation of 76 TF-encoding genes (58 upregulated, 18 downregulated) was found reproducibly in at least three different experiments in *ap2c1 mkp1* double mutant plants. Among them, genes encoding members of the WRKY family were most abundant: 15 *WRKY* genes were upregulated (Figure 6, Supplementary Table IV) and one downregulated (Supplementary Table IV). A further prevalent group of TF-encoding genes affected in *ap2c1 mkp1* plants includes *AP2*/*ERF* described for their involvement in development, including *RAP2.6L* (Yang *et al*., 2018) and *WIND3* (Smit *et al*., 2020) (Figure 7). ANAC TF family members are implicated in senescence and stress-related processes (Bu *et al*., 2008; Jensen *et al*., 2010; Saga *et al*., 2012; Wu *et al*., 2012). Our results show that several ANAC TF-encoding genes are upregulated in *ap2c1 mkp1* plants (Figure 8). Thus, our data suggest a cooperative function of AP2C1 and MKP1 in the transcriptional regulation of a set of *WRKY, AP2*/*ERF* and *ANAC* genes in the WT.

**Figure 5.**
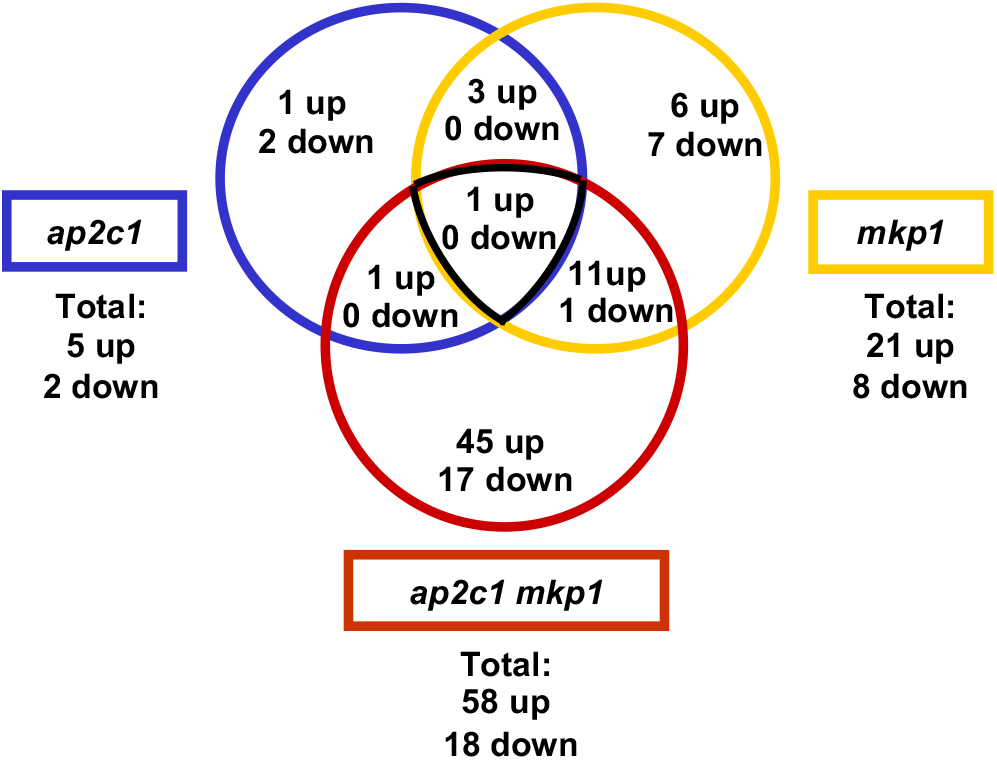
Venn diagram of TFs differentially expressed in MAPK phosphatase mutant plants. The number of genes at least three-fold up- or downregulated in *ap2c1*, in *mkp1* and in *ap2c1 mkp1* plants compared to WT in three biological replicates is indicated. The expression of 1,880 TF-encoding genes was analysed.

**Figure 6.**
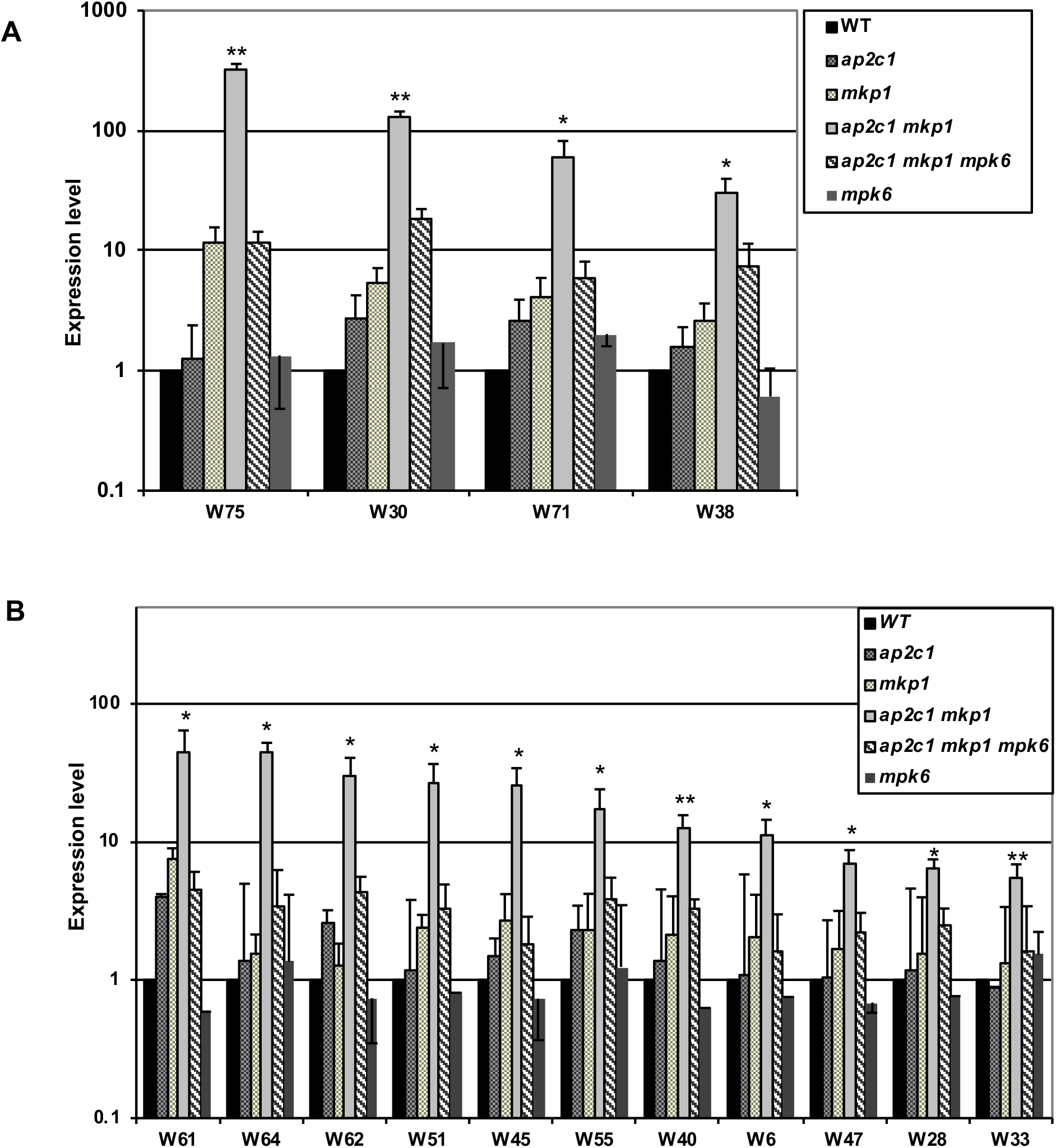
Expression of WRKY-encoding (W) genes. The transcript levels of WRKY-encoding (W) genes were quantified by RT-qPCR in *ap2c1, mkp1, ap2c1 mkp1, ap2c1 mkp1 mpk6*, and *mpk6* plants, and compared to WT (values set to 1). Bars represent mean values of three replicates ± SD. **A.** Upregulation of *WRKY75*, *WRKY30*, *WRKY71*, and *WRKY38* transcript levels in *ap2c1 mkp1* is not solely dependent on MPK6. Data are expressed on a log_10_ scale after normalisation over WT values. **B.** Upregulation of *WRKY61*, *WRKY64*, *WRKY62*, *WRKY51*, *WRKY45*, *WRKY55*, *WRKY40*, *WRKY6*, *WRKY47*, *WRKY28* and *WRKY33* transcript levels depends on MPK6, *p < 0.05, **p < 0.01 Mann-Whitney *U* test.

**Figure 7.**
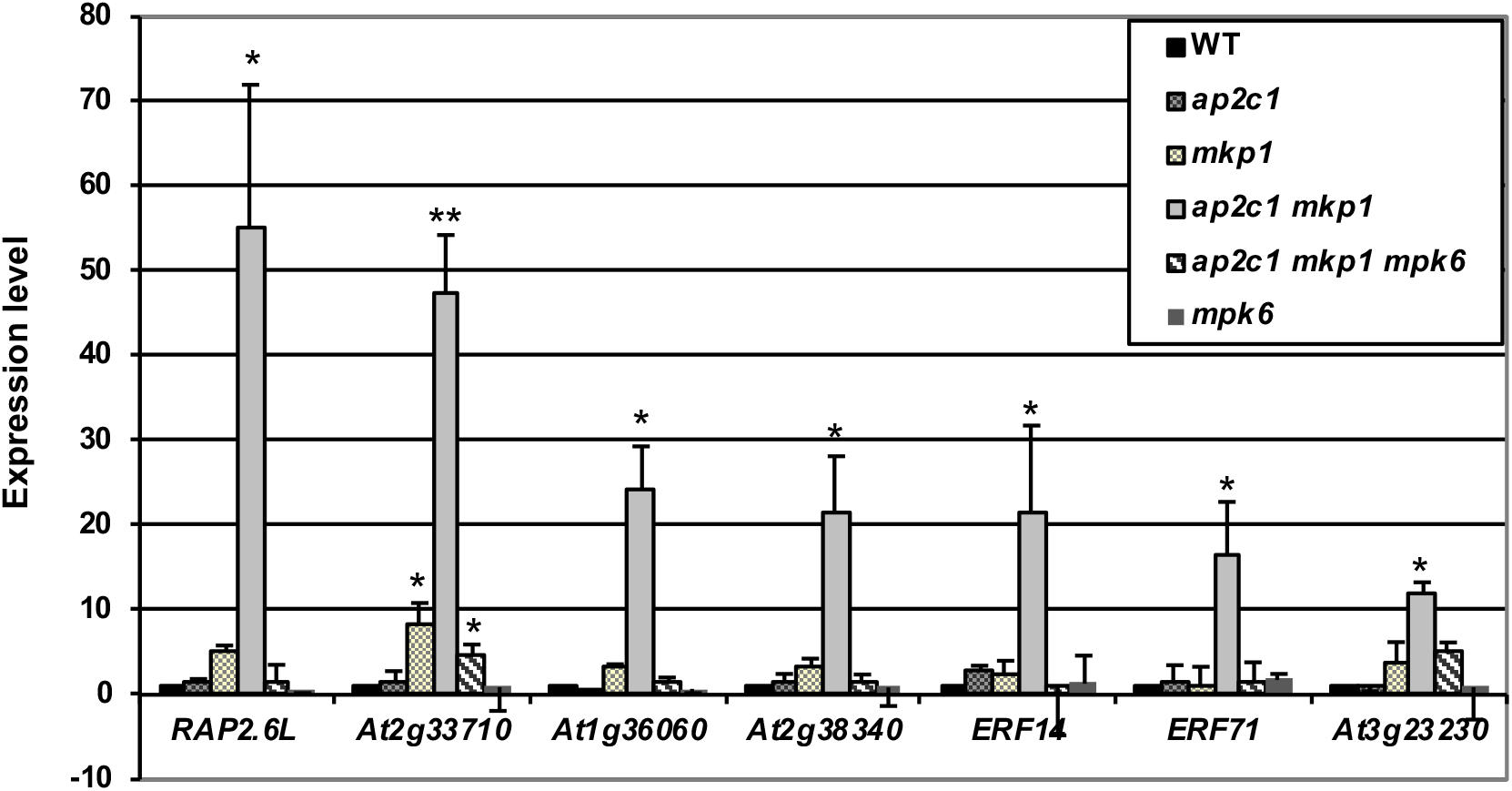
Genes encoding members of the AP2/ERF TF family are highly upregulated in *ap2c1 mkp1* plants. Transcript levels of AP2/ERF-encoding genes were quantified by RT-qPCR in *ap2c1*, *mkp1*, *ap2c1 mkp1*, *ap2c1 mkp1 mpk6*, and *mpk6* plants and compared to WT, where expression levels were set to 1. Bars represent mean values of at least three replicates ± SD. Strong upregulation of *At2g33710*, *At1g71520* and *At3g23230* (*ERF98*/*TDR1*) in *ap2c1 mkp1* is not solely dependent on MPK6, while upregulation of *RAP2.6L*, *WIND3*, *ERF1*, *DREB19*, *ERF14*, and *ERF17* in *ap2c1 mkp1* is almost completely dependent on MPK6, *p < 0.05, **p < 0.01 Mann-Whitney *U* test.

**Figure 8.**
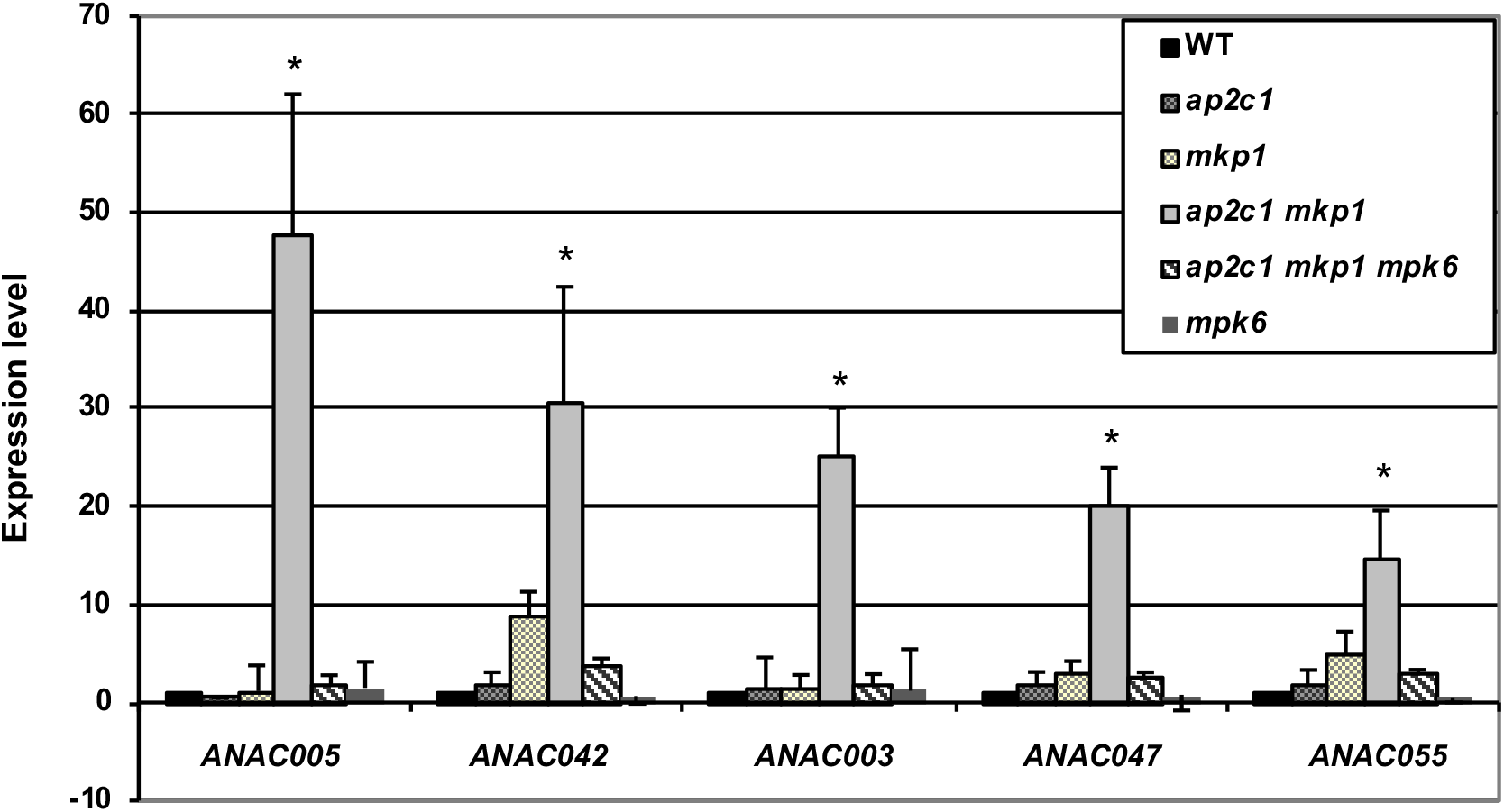
Genes encoding members of the ANAC TF family are highly upregulated in *ap2c1 mkp1* plants. Transcript levels were quantified by RT-qPCR in *ap2c1, mkp1, ap2c1 mkp1, ap2c1 mkp1 mpk6*, and *mpk6* plants and compared to WT, where expression levels were set to 1. Bars represent mean values of at least three replicates ± SD. *ANAC005*, *ANAC042*, *ANAC003*, *ANAC047*, and *ANAC055* are upregulated in *ap2c1 mkp1* in a MPK6-dependent manner, while in *ap2c1 mkp1 mpk6* plants their levels are similar to WT, *p < 0.05, Mann-Whitney *U* test.

Our observation that *ap2c1 mkp1 mpk6* plants are phenotypically much less affected than *ap2c1 mkp1* double mutants suggested that severe phenotypic aberrations in the latter are mediated by MPK6. This prompted us to investigate *ap2c1 mkp1 mpk6* plants for the expression of TFs misregulated in *ap2c1 mkp1*. Indeed, 50 of the 76 TF genes strongly affected in *ap2c1 mkp1* plants were not altered in their expression in *ap2c1 mkp1 mpk6*, linking MPK6 over-activation to their misexpression. Among these, expression of *WRKY6, WRKY28, WRKY33* and *WRKY45* (Figure 6B), the AP2/ERF family members *RAP2.6L*, *WIND3*, *ERF1*, *DREB19*, *ERF14*, *and ERF17* (Figure 7), and of several ANAC TF-encoding genes, such as *ANAC005*, *ANAC042*, *ANAC003*, *ANAC047*, *and ANAC055* (Figure 8) are dependent on the presence of MPK6. The expression of *WRKY75*, *WRKY71*, *WRKY38* and *WRKY30* (Figure 6), and of the *AP2/ERF* genes *At2g33710, At1g71520* and *At3g23230* remained upregulated more than 5-fold in the absence of MPK6 in *ap2c1 mkp1 mpk6* plants (Figure 7), suggesting that regulation of these TFs is controlled by other factors (possibly other MAPKs).

### Defence responses, camalexin, SA and the senescence marker gene *SENESCENCE-ASSOCIATED GENE12* (*SAG12*) are upregulated in *ap2c1 mkp1* plants

It has been shown previously that *mkp1* plants accumulate higher levels of the phytoalexin camalexin (Bartels *et al*., 2009). To investigate if the expression of genes encoding camalexin biosynthesis enzymes was affected in *ap2c1 mkp1* plants, we studied the expression of a key gene in the pathway, *CYP71B15*/*PAD3*. A strong upregulation (more than 300-fold, respectively) was detected in *ap2c1 mkp1* plants compared to WT (Figure 9). Moreover, *ap2c1 mkp1 mpk6* plants still had remarkably high transcript levels (upregulation ca. 10-fold) of *CYP71B15*/*PAD3*, showing that MPK6 is an important, but not the sole factor behind its upregulation in *ap2c1 mkp1* plants. Also, *mkp1* single mutant plants showed a >10-fold upregulation of the gene (Figure 9A).

**Figure 9.**
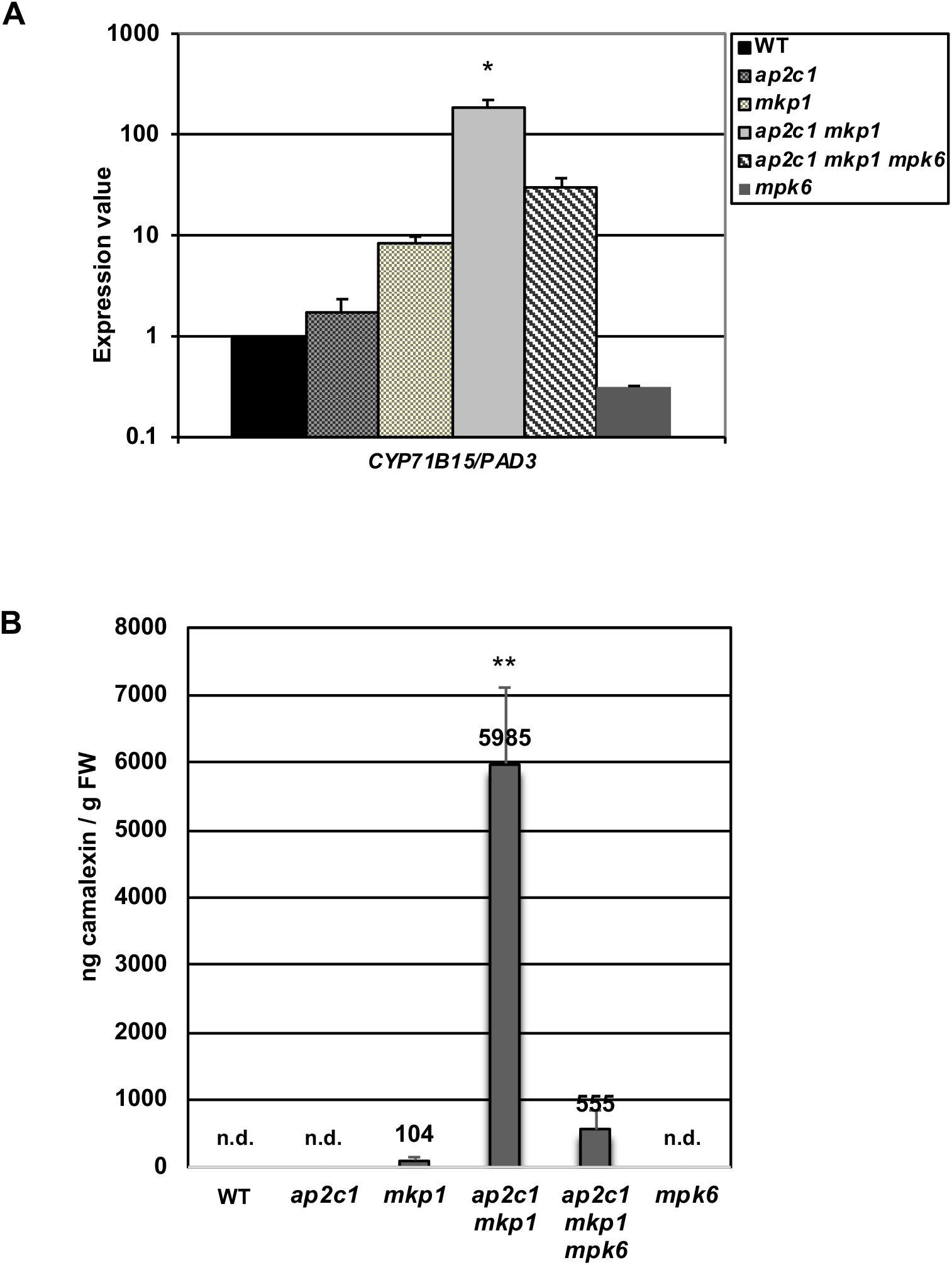
Upregulation of camalexin biosynthetic gene *CYP71B15*/*PAD3* and camalexin accumulation in *ap2c1 mkp1* plants are mostly mediated by MPK6. **A.** Transcript level of *CYP71B15*/*PAD3*, an enzyme required for camalexin biosynthesis, was quantified by RT-qPCR in *ap2c1, mkp1, ap2c1 mkp1, ap2c1 mkp1 mpk6*, and *mpk6* plants and compared to WT, where expression levels were set to 1. Bars represent mean values of at least three replicates ± SD, expressed on a log_10_ scale. **B.** Levels of total camalexin determined by HPLC in leaves of 4-week-old WT, *ap2c1*, *mkp1*, *ap2c1 mkp1*, *ap2c1 mkp1 mpk6* and *mpk6* plants. Results shown are mean with SE (n=4), n.d. = not detected, *p < 0.05, **p < 0.01, Student’s *t*-test.

To investigate if the increased *CYP71B15*/*PAD3* expression level correlates with camalexin accumulation, total camalexin was quantified in WT and mutant plants. Indeed, in agreement with previous findings (Bartels *et al*., 2009) we found increased camalexin levels in *mkp1* plants and very high camalexin accumulation in the *ap2c1 mkp1* mutant, which was not solely dependent on MPK6 (Figure 9B).

Upregulation of MAPK activities and macroscopic lesion formation in leaves of *ap2c1 mkp1* indicated the possible activation of a hypersensitive-like response in these plants. Since this is associated with the accumulation of the stress hormone SA, we measured SA in leaves of *ap2c1 mkp1, ap2c1 mkp1 mpk6* as well as in WT and single mutants. Indeed, we found a 35-fold increase of SA in *ap2c1 mkp1* plants compared to WT (Figure 10), whereas *ap2c1, ap2c1 mkp1 mpk6*, and *mpk6* plants showed SA amounts similar to the WT. In agreement with previous data (Bartels *et al*., 2009), we detected enhanced total SA amounts (>2-fold) also in *mkp1* plants compared to WT (Figure 10).

**Figure 10.**
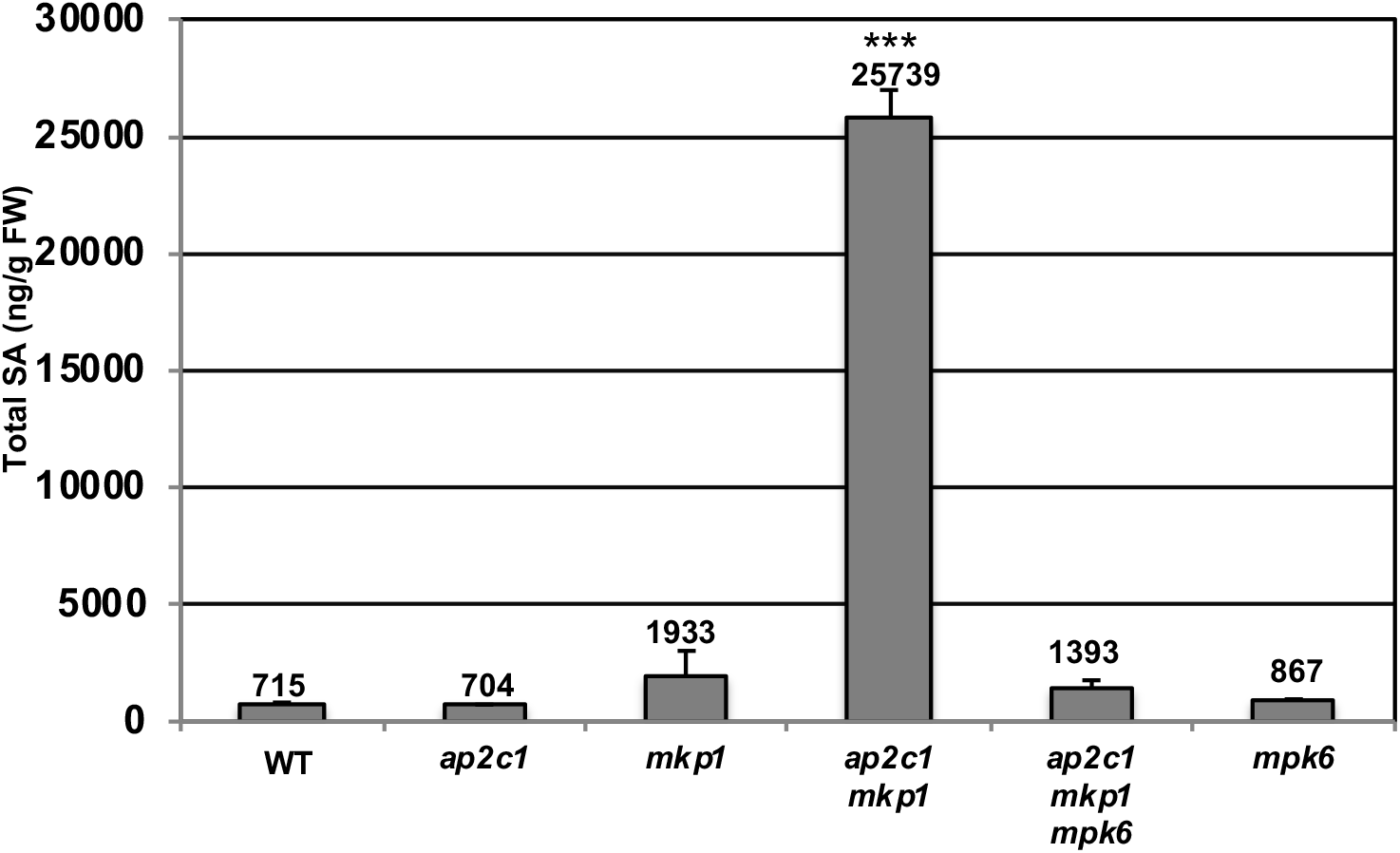
*ap2c1 mkp1* plants accumulate high levels of SA in a MPK6-dependent manner. Total SA levels of five-week-old WT, *ap2c1*, *mkp1*, *ap2c1 mkp1*, *ap2c1 mkp1 mpk6*, and *mpk6* plants grown in standard short-day conditions, determined by HPLC and expressed as ng per g FW. Error bars represent SD of four biological replicates, ***p < 0.001, Student’s *t*-test.

Leaf necrosis observed in *ap2c1 mkp1* leaves (Figure 1B) and the upregulation of *WRKY6* (Figure 6B), which is a senescence-related marker gene (Rushton *et al*., 2010), suggested that early senescence was induced in these plants. Thus, we investigated the expression of the senescence-specific marker gene *SAG12* (Noh and Amasino, 1999) and found that it strongly upregulated in *ap2c1 mkp1* plants, dependent on MPK6 (Figure 11). This data along with the upregulation of *WRKY6* (Figure 6B) indicates aberrant, early induction of senescence-related processes in the double phosphatase mutant.

**Figure 11.**
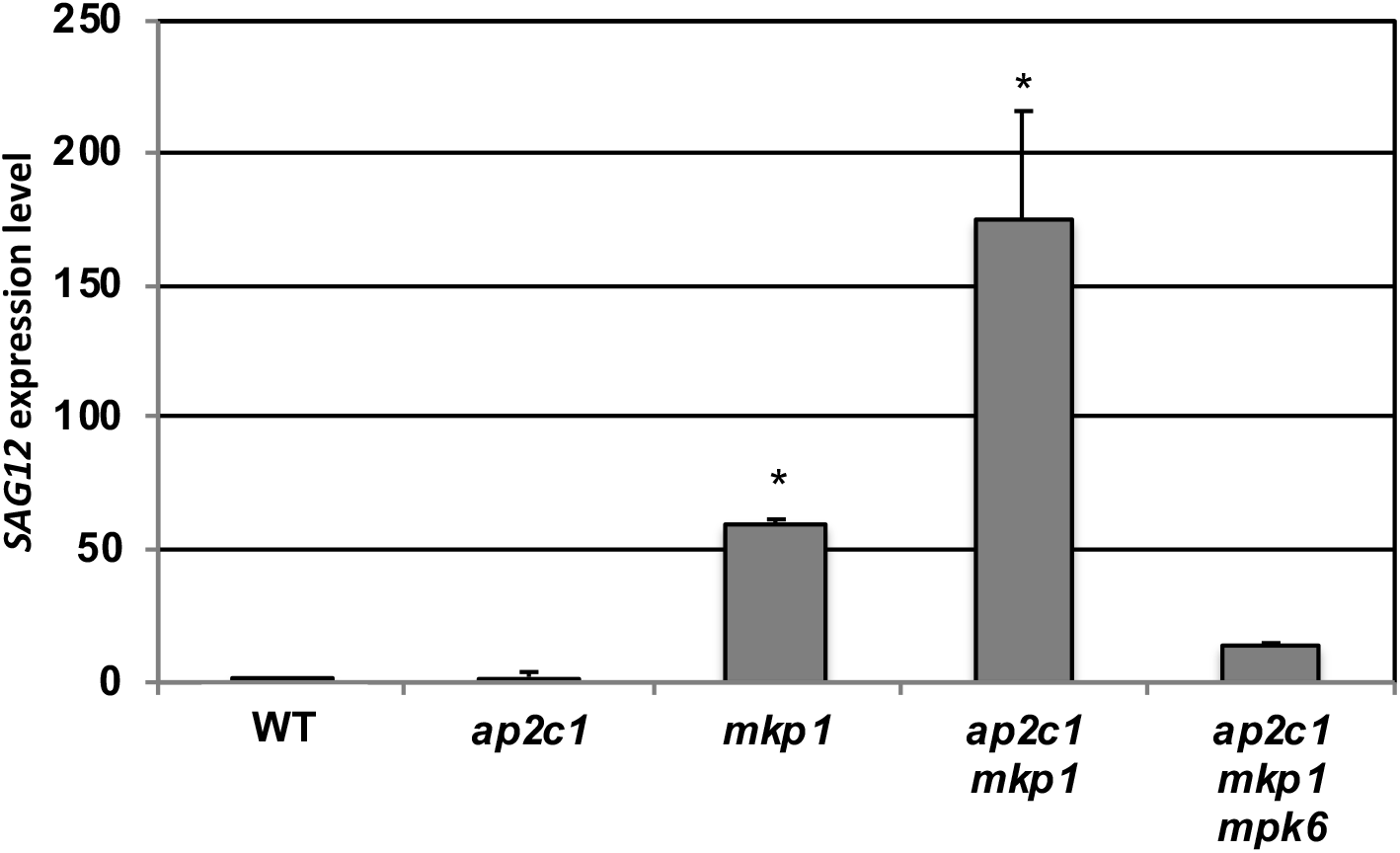
Upregulation of the senescence-marker gene *SAG12* in *ap2c1 mkp1* plants is mainly mediated by MPK6. RT-qPCR quantification of *SAG12* transcript level in leaves of six-week-old *ap2c1*, *mkp1*, *ap2c1 mkp1*, and *ap2c1 mkp1 mpk6* mutant plants compared to WT plants grown in standard short-day conditions. The *ap2c1 mkp1* mutant displays more than 160-times higher *SAG12* transcript level than WT. Of note, *SAG12* upregulation was attenuated in the *ap2c1 mkp1 mpk6* triple mutant (14-times upregulation compared to WT). Error bars represent SD of three biological replicates, *p < 0.05, Student’s *t*-test.

## Discussion

### Coordinated control of MAPK activities by AP2C1 and MKP1

Acclimation for survival is a fundamental principle, which relies on intracellular signalling in every organism. Different signals converge at the level of MAPK cascades, and from there diverge into a range of different downstream pathways and responses (Andreasson and Ellis, 2010; Rasmussen *et al*., 2012; Rodriguez *et al*., 2010). Considering the broad spectrum of signals transduced by overlapping players of MAPK pathways it is puzzling how response specificity is attained (Lampard *et al*., 2009; Meng and Zhang, 2013; Rodriguez *et al*., 2010). Several signalling scenarios have been investigated that could help explain pathway specificity, including activity-dependent kinase distribution and localization, protein complex formation (e.g. interaction with scaffolding proteins), and dephosphorylation by protein phosphatases (Krysan and Colcombet, 2018). Over the last decades mainly the functions of MPK3/MPK4/MPK6 in diverse pathways have been described, indicating them as both points of divergence and integration hubs in cellular signalling (Bigeard and Hirt, 2018; Peng *et al*., 2018).

Here, we provide evidence that two evolutionarily distinct MAPK phosphatases control stress-related signalling in Arabidopsis by inactivating an overlapping set of target MAPKs that mediate stress and defence responses. The Ser/Thr PP2C phosphatase AP2C1 and the dual-specificity phosphatase MKP1 contribute to ensure appropriate inactivation of MAPKs during stress. Both AP2C1 and MKP1 target MPK3, MPK4 and MPK6 (Anderson *et al*., 2011; Bartels *et al*., 2009; Galletti *et al*., 2011; Schweighofer *et al*., 2007; Shubchynskyy *et al*., 2017; Sidonskaya *et al*., 2016; Ulm *et al*., 2002). Enhanced activation of MAPKs by wounding and constitutive stress signalling in the absence of stress in *ap2c1 mkp1* plants indicate that the lack of both MAPK phosphatases creates a shortfall downstream of MAPKs, exemplified by deregulated expression of TF-encoding genes.

An enhanced kinase activity in *mkp1* plants *versus* WT at the earlier time points after wounding compared to *ap2c1 versus* WT suggests that the contribution of MKP1 to inactivating MAPKs is already set before wounding, or during a very early stage of signalling. On the contrary, AP2C1 adds to MAPK inactivation at later time points. It is possible that AP2C1 is primarily responsible for keeping the stress-induced activation below a certain threshold and controlling the duration of kinase activation during acute stress acting as an “emergency brake”, while MKP1 is predominantly responsible for suppressing kinase activities under normal conditions, providing a “constitutive brake”. This hypothesis is supported by the demonstrated induction of *AP2C1* expression by a plethora of stresses, while *MKP1* shows comparatively marginal changes in expression (https://www.genevestigator.com). These observations are also consistent with a recent comprehensive analysis of the Arabidopsis proteome, which covers more than 14,000 proteins and where in ambient conditions the overall MKP1 abundance outnumbers by far that of AP2C1 (http://athena.proteomics.wzw.tum.de/) (Mergner *et al*., 2020), underlining the rather specific role of AP2C1 under stress conditions. The AP2C1 paralogues AP2C2 and AP2C3 (Schweighofer *et al*., 2014; Umbrasaite *et al*., 2010; Umbrasaite *et al*., 2011) as well as MKP1 and PTP1 interact with the same MAPKs and dephosphorylate them to various extents (Bartels *et al*., 2009). However, the rather mild phenotypes of *ap2c2 mkp1* and *ap2c3 mkp1* plants and the WT-like appearance of *ap2c1 ptp1* (this work) compared to *ap2c1 mkp1* plants clearly indicate specific genetic interactions and redundant functions of the evolutionary distant AP2C1 and MKP1 phosphatases in the regulation of signalling pathways.

### Manifestation of cell death in *ap2c1 mkp1* plants

The lesions in leaves of *ap2c1 mkp1* plants suggest autoimmune-like responses most likely caused by misregulation of MAPKs and/or failed control of guarding resistance (R) proteins (Rodriguez *et al*., 2016). AP2C1 and MKP1 share the target MAPKs MPK3, MPK4 and MPK6, where MPK4 and some of its upstream MAPK cascade members were originally described as negative regulators of plant immunity based on their mutant plant phenotypes, for example MEKK1 and MKK1/2 (Petersen *et al*., 2000; Rasmussen *et al*., 2012). The improper activation of the *R*-gene *SUMM2* is mainly responsible for the phenotypical defects of the *mpk4* mutant and of other mutant plants in the pathway, identifying the MEKK1-MKK1/2-MPK4 module as a positive regulator of stress responses (Zhang *et al*., 2012). Similar observations connecting phosphatase-targeted MAPKs with autoimmune-like phenotypes have been made by ectopically expressing constitutively active MPK3 (Genot *et al*., 2017) or by inducibly expressing MKK5, which activates MPK3 and MPK6 (Lassowskat *et al*., 2014). Both approaches led to a plethora of phenotypic and molecular changes including dwarfism, lesion formation, de-repression of defence gene expression, and the accumulation of stress hormones, similar to the *ap2c1 mkp1*-related phenotypes described in this work (see Results).

The single *mkp1* and the double *mkp1 ptp1* mutants show constitutive defence responses including increased levels of SA and camalexin, suggesting partially overlapping functions of MKP1 and PTP1 in repressing SA biosynthesis (Bartels *et al*., 2009). Similarly, the strong accumulation of SA and camalexin in *ap2c1 mkp1* compared to *mkp1* plants suggests a collaborative action of both AP2C1 and MKP1 as negative regulators of SA and camalexin production (this work). This accumulation is probably MPK6-dependent, as the introduction of the *mpk6* mutation in *ap2c1 mkp1 mpk6* plants restores SA and camalexin levels similar to those of WT and *mkp1 ptp1 mpk6* mutant (Bartels *et al*., 2009). Notably, rescue of the severe *ap2c1 mkp1* growth phenotypes by elevated temperature is in accordance with the observed temperature dependency of SA-related phenotypes (Ichimura *et al*., 2006; Su *et al*., 2007; Suarez-Rodriguez *et al*., 2007), as well as with the suppression of *SNC1* expression and reduction of SNC1 activity by high temperature (Yang and Hua, 2004; Zhu *et al*., 2010). The resistance protein SNC1 is a modifier of *mkp1* in the Col-0 accession, where partial rescue of *mkp1* and *mkp1 ptp1* growth phenotypes by a loss-of-function *snc1* mutation indicates a sensitized SNC1 signaling pathway in the absence of MKP1 (Bartels *et al*., 2009).

Previous findings that SA acts together with ET to regulate cell death (Rao *et al*., 2002), the requirement of ET biosynthesis for H_2_O_2_ accumulation and subsequent cell death (Overmyer *et al*., 2003), and the induction of cell death in Arabidopsis leaves by persistent activation of MAPKs with gain-of-function MKK4 and MKK5 (Ren *et al*., 2002) all correlate with the cell death phenotype observed in the *ap2c1 mkp1* mutant, where MAPKs - and other stress-related factors - may be (hyper)-activated. Therefore, we conclude that the majority of the phenotypes observed in *ap2c1 mkp1* plants, both visible and molecular, are due to the misregulation of MAPK pathways, even in the absence of stress.

### AP2C1 and MKP1 affect MAPK-regulated ET biosynthesis

Activated MPK6 controls ET levels by both inducing the transcription of *ACS* family genes and by phosphorylating ACS proteins, the rate-limiting enzymes in ET biosynthesis. Phosphorylated ACSs become more stable and, thus, ET synthesis is enhanced by elevated MPK6 activity (Kim *et al*., 2003; Li *et al*., 2012; Liu and Zhang, 2004; Xu *et al*., 2008). In *ap2c1 mkp1*, the enhanced ET production is certainly due to, at least in part, the highly increased expression of *ACS6* compared to WT. A considerable additive effect on ET overproduction by the double *ap2c1 mkp1* mutation suggests that even though MKP1 is a determining MAPK phosphatase affecting ET production, there are overlapping and non-redundant functions of AP2C1 and MKP1 in the regulation of stress-induced ET biosynthesis. Detection of enhanced and MPK6-dependent expression of *WRKY33*, encoding a TF that binds to the promoter of *ACS* genes and is a substrate of MPK3/MPK6, suggests an involvement of WKRY33 itself in *ACS* overexpression in *ap2c1 mkp1* plants (this work and (Li *et al*., 2012)). The identification of genes encoding TFs of the AP2/ERF family members (ET-responsive element-binding proteins) among the uppermost induced ones in *ap2c1 mkp1* plants suggests a path to enhanced ET amounts in these plants.

### AP2C1 and *MKP1* control the expression of stress-responsive TF-encoding genes, predominantly *via* MPK6

Transcriptional reprogramming in response to activated MAPK signalling suggests an involvement of TFs. Our results indicate that the concomitant lack of the MAPK regulators AP2C1 and MKP1 results in elevated basal MAPK activities and leads to highly enhanced expression of *WRKY* TF genes, in some cases by more than hundred-fold compared to WT. The *ap2c1 mkp1* mutant phenotypes and the described functions of some upregulated *WRKYs* indicate that stress responses are constitutively active in these plants. This correlates with reports demonstrating an involvement of WRKYs in oxidative stress responses, in the induction of ET and camalexin biosynthesis (*WRKY30, WRKY33*), in the response to pathogens (*WRKY71, WRKY40*), in basal defence (*WRKY38, WRKY42*), and defence- and senescence-related processes (*WRKY6*) (Rushton *et al*., 2010).

Direct feedback mechanisms among WRKYs themselves have been shown (Mao *et al*., 2011) and are generally proposed, where WRKYs positively auto-regulate their own gene expression and/or cross-regulate expression of other *WRKY* genes (Birkenbihl *et al*., 2017; Mao *et al*., 2011; Pandey and Somssich, 2009). Thus, it could be that the enhanced activation of MAPKs in *ap2c1 mkp1* plants leads to phosphorylation and thus activation of MAPK target WRKY proteins, which serve as activated TFs for a further series of *WRKY* genes. In any case, MPK6 seems to be a major player responsible for mediating the upregulation of several *WRKYs, AP2*/*ERFs, ANACs* and other TF-encoding genes. MPK6 controls the expression of several *WRKYs* to different extents, as shown in *ap2c1 mkp1* plants compared to *ap2c1 mkp1 mpk6* (Figure 6A). This data demonstrates that not only MPK6 but also other factor(s) affect *WRKY* gene expression. MPK3, as the closest paralogue of MPK6, may be a possible candidate, since MPK3 is also more strongly activated in *ap2c1 mkp1* than in the respective single mutant plants. We confirmed MPK6-dependent *WRKY33* expression (Mao *et al*., 2011); however, the higher MPK4 activities in *ap2c1 mkp1* may also lead to higher amounts of active WRKY33 protein (Birkenbihl *et al*., 2017; Qiu *et al*., 2008). Thus, our data suggest that AP2C1 and MKP1 may play a dual role in regulating camalexin biosynthesis, on the one hand by controlling MPK6 activity, which positively regulates *WKRY33* expression, and on the other by controlling MPK4 activity, which in turn stimulates WRKY33 leading to transactivation of *CYP71B15/PAD3*.

A PAMP-activated MPK3/MPK6 pathway was previously reported to elevate *WRKY22* and *WRKY29* expression (Asai *et al*., 2002). Strongly enhanced MPK3/MPK6 activities, but unaffected expression of either *WRKY22* or *WRKY29* in untreated *ap2c1 mkp1* plants, show that for *WRKY22*/*29* overexpression the MPK6 hyperactivation is sufficient (Asai *et al*., 2002) but not necessary (this work) and that other factors (possibly MAPKs) may be playing a role instead of MPK6.

### Senescence is *repressed* by AP2C1 and MKP1 phosphatases in an MPK6-dependent way

Several lines of evidence indicate that the *ap2c1 mkp1* mutant undergoes precocious senescence. Leaf senescence is a highly regulated process that finally leads to cell death and tissue disintegration, at the same time contributing to the fitness of the whole plant. Senescence is controlled by endogenous and environmental cues, and can be triggered prematurely by different abiotic/biotic stresses due to pathogen attack, wounding, UV light irradiation, and high ozone levels (Hanfrey *et al*., 1996; He *et al*., 2001; John *et al*., 2001; Lim *et al*., 2007; Miller *et al*., 1999). The MKK9-MPK6 cascade has been shown to positively regulate leaf senescence in Arabidopsis (Zhou *et al*., 2009). Hyperactivation of MPK6 and other MAPKs, in addition to autoimmune-like responses, also promotes senescence, which is very evident in older leaves of *ap2c1 mkp1* plants and correlates with significant upregulation of the senescence-specific marker gene *SAG12* (Guo and Gan, 2005; Noh and Amasino, 1999). Partial suppression of *SAG12* overexpression in *ap2c1 mkp1 mpk6* suggests an MPK6-dependent regulation (possibly involving other MAPKs) in promoting plant senescence.

Genome-wide transcriptomics previously identified several senescence-related TFs from the ANAC family (Breeze *et al*., 2011). We could highlight strong MPK6-dependent induction of *ANAC005*, *JUB1*/*ANAC042* (Saga *et al*., 2012; Shahnejat-Bushehri *et al*., 2016; Wu *et al*., 2012), *ANAC003*/*XVP* (Yang *et al*., 2020), *ANAC047* (Mito *et al*., 2011), and *ANAC055* (Bu *et al*., 2008; Hickman *et al*., 2013; Schweizer *et al*., 2013; Tran *et al*., 2004) in *ap2c1 mkp1* plants. This induction of senescence-related TFs reveals a novel link between senescence-related processes and MAPK signalling.

We conclude that the induction of senescence processes as well as hypersensitive response-like cell death results in premature death of leaves in *ap2c1 mkp1* plants. The crosstalk between senescence and abiotic stress or pathogen responses is accentuated in *ap2c1 mkp1* plants where upregulation of TFs involved in these processes is happening.

Taken together, our results show that two evolutionarily unrelated MAPK phosphatases, AP2C1 and MKP1, perform both distinct and overlapping functions in the regulation of stress-induced MPK3, MPK4 and MPK6 activities. Our genetic dissection indicates that the known role of MPK6 in mediating cell death, ET-, SA- and senescence-related phenotypes is attenuated by both AP2C1 and MKP1. It also demonstrates that the expression of specific TF-encoding genes is affected by MAPK(s) hyperactivation due to the lack of these two MAPK phosphatases *in planta*, revealing potential new target genes downstream of MPK6 signalling. Additionally, our data suggest new roles for MPK3 or MPK4 in the regulation of cell signalling. In the future, the study of individual and combinatorial mutants will allow us to genetically disentangle the contribution of specific protein kinases and phosphatases to complex signalling networks and downstream cell responses.

## Supplementary data

**Supplementary Figure 1:** Phenotypes of Arabidopsis single, double and triple mutant plants.

**Supplementary Figure 2:** Loss of both AP2C1 and MKP1 leads to severe phenotypes in growth and development, which are mediated by MPK6.

**Supplementary Table I.** Expression of TF-encoding genes modulated by the absence of AP2C1 but not MKP1.

**Supplementary Table II.** Expression of TF-encoding genes modulated by the absence of MKP1 but not of AP2C1.

**Supplementary Table III.** Expression of TF-encoding genes modulated by the absence of both, MKP1 and AP2C1.

**Supplementary Table IV.** TF-encoding genes deregulated in *ap2c1 mkp1* plants.

## Acknowledgements

We thank Verena Ibl (University of Vienna, AT), Mary G. Wallis (University of Applied Sciences, FH Campus Vienna, AT) and Francesca Cardinale (University of Turin, IT) for critical reading of the manuscript, and the Nottingham Arabidopsis Stock Centre for providing SALK mutant lines. This work was supported by the Austrian Science Fund (FWF) with the grants I-255, W1220-B09 to IM and P27254 to IM/AS. SeB was supported by the German Research Foundation (DFG) through the Research Training Group GRK1305. BMR and SaB thank the MPI of Molecular Plant Physiology (Potsdam, DE) for financial support.

## Author contributions

ZA, VK, VS, KK, MaS, WR, MiS, SeB, SaB and AS performed experiments, ZA, VK, VS, KK, FM, RU, SaB, BMR, IM and AS designed experiments; and ZA, SaB, BMR, IM and AS wrote the paper.

## Dedication

The authors dedicate this article to the memories of:

Irute Meskiene (1956-2017)

(Paškauskas *et al*., 2017)

Manfred Schwanninger (1963-2013)

(Meder, 2014)

## Data availability statement

The data supporting the findings of this study are available from the corresponding author (Alois Schweighofer), upon request.

## Supplementary information

**Supplementary Table I. Expression of TF-encoding genes modulated by the absence of AP2C1 but not MKP1.** Transcript levels were quantified by RT-qPCR and expressed as log_2_ of fold change (FC) for each mutant compared to WT. Genes affected by at least 1.56-fold only in *ap2c1* while not more than 0.9-fold in *mkp1* are listed. Data are from two independent biological replicates and are reported with SE.

**Supplementary Table II. Expression of TF-encoding genes modulated by the absence of MKP1 but not of AP2C1.** Transcript levels were quantified by RT-qPCR and are expressed log_2_ of fold change (FC) for each mutant compared to WT. Genes affected by at least 1.56-fold only in *mkp1* while not more than 0.9-fold in *ap2c1* are listed. Data are from two independent biological replicates and are reported with SE.

**Supplementary Table III. Expression of TF-encoding genes modulated by the absence of both MKP1 and AP2C1.** Transcript levels were quantified by RT-qPCR and are expressed as fold change (FC) in log_2_ scale for each of the mutants compared to WT. Genes affected by at least 1.56-fold both in *ap2c1* and *mkp1* are listed. Data are from two independent biological replicates and are reported with SE.

**Supplementary Table IV. TF-encoding genes deregulated in *ap2c1 mkp1* plants.** Gene transcript levels were quantified by RT-qPCR and expressed as fold change in log_2_ scale for each of the mutants compared to WT. Genes affected in *ap2c1 mkp1* compared to WT by at least 1.56-fold in log_2_ scale are listed. Data are from three independent biological replicates and are reported with SD.

**Supplementary Figure 1. Phenotypes of Arabidopsis single, double and triple mutant plants.**

**A.** Phenotypes of WT, *ap2c1*, and *mkp1* plants grown for three weeks in long-day conditions. **B.** Phenotypes of WT, *ap2c1 mkp1, ap2c2 mkp1*, and *ap2c3 mkp1* plants grown for four weeks in short-day conditions. **C.** Phenotypes of WT, *ap2c1, ap2c2, ap2c3, mkp1, ap2c1 mkp1, ap2c2 mkp1, ap2c3 mkp1, mpk6*, and *ap2c1 mkp1 mpk6* plants grown for four weeks in short-day condition, followed by three weeks in long-day condition.

**Supplementary Figure 2. Loss of both AP2C1 and MKP1 leads to severe phenotypes in growth and development, which are mediated by MPK6.**

**A.** Phenotypes of WT (left) and *ap2c1 mkp1* (right, indicated by arrow) grown for 2.5 weeks in standard long-day conditions. During the first 18 days of growth, the phenotypic differences of *ap2c1 mkp1* compared to WT plants are visibly manifested as a difference in plant size. **B.** After approximately 3.5 weeks (26 days), premature death of leaf tissue as well as abnormal leaf growth and morphology in *ap2c1 mkp1* plants (right, indicated by arrow) became apparent. Scale bars in **A** and **B** = 1 cm. **C.** Close-up of *ap2c1 mkp1* plant grown for 3.5 weeks in standard long-day condition. **D.** Phenotype of seven-week-old *ap2c1 mkp1* plant grown in standard long-day condition. **E.** Close-up of seven-week-old *ap2c1 mkp1* plant showing misshaped inflorescence. **F, G.** Phenotypes of eight-week-old WT, *ap2c1*, *mkp1*, *ap2c1 mkp1*, and *ap2c1 mkp1 mpk6* plants grown for the first six weeks in short-day and for a further two weeks in long-day conditions. The *ap2c1 mkp1* double mutant displays a severe dwarf phenotype, premature leaf decay, lack of normal shoot development, and strongly impaired inflorescence growth. The inset picture above shows a close-up of the *ap2c1 mkp1* plant shown. In *ap2c1 mkp1 mpk6* triple mutant plants these phenotypes were rescued.

