## Supplementary Material for "Dual control of MAPK activities by AP2C1 and MKP1 MAPK phosphatases regulates defence responses in Arabidopsis"

**Supplementary Table I. Expression of TF-encoding genes modulated by absence of AP2C1 but not of MKP1.** Transcript levels were quantified by RT-qPCR and expressed as $log_2$ of fold change (FC) for each mutant compared to WT. Genes affected by at least 1.56 only in *ap2c1* while not more than 0.9 in *mkp1* are listed. Data are from 2 independent biological replicates and are reported with SE.

| AGI | gene family | gene name(s) | *ap2c1* $log_2$ FC | SE of -ΔΔct | *mkp1* $log_2$ FC | SE of -ΔΔct |
|---|---|---|---|---|---|---|
| At2g46130 | WRKY | WRKY43 | **1.56** | 0.40 | -0.46 | 0.77 |
| At1g65130 | C2H2 | | **-1.90** | 0.13 | -0.42 | 0.29 |
| At3g27810 | MYB | MYB21/MYB3 | **-3.79** | 0.32 | -0.27 | 0.35 |

**Supplementary Table II. Expression of TF-encoding genes modulated by absence of MKP1 but not of AP2C1.** Transcript levels were quantified by RT-qPCR and are expressed $log_2$ of fold change (FC) for each mutant compared to WT. Genes affected by at least 1.56 only in *mkp1* while not more than 0.9 in *ap2c1* are listed. Data are from 2 independent biological replicates and are reported with SE.

| AGI | gene family | gene name(s) | *ap2c1* $log_2$ FC | SE of -ΔΔct | *mkp1* $log_2$ FC | SE of -ΔΔct |
|---|---|---|---|---|---|---|
| At5g43650 | bHLH | BHLH92 | -0.19 | 0.75 | **3.77** | 0.27 |
| At1g71520 | AP2/EREBP | | 0.41 | 0.91 | **3.71** | 0.54 |
| At5g13080 | WRKY | WRKY75 | 0.61 | 0.13 | **3.21** | 0.09 |
| At3g49760 | bZIP | BZIP5 | -0.40 | 0.69 | **2.49** | 0.20 |
| At2g43000 | NAC | ANAC042/JUB1 | 0.61 | 0.04 | **2.36** | 0.29 |
| At2g33710 | AP2/EREBP | | 0.34 | 0.10 | **2.29** | 0.35 |
| At5g56960 | bHLH | | -0.14 | 0.25 | **2.29** | 0.15 |
| At2g47190 | MYB | MYB2 | 0.48 | 0.11 | **2.23** | 0.30 |
| At1g66600 | WRKY | WRKY63 | -0.60 | 0.25 | **2.13** | 0.70 |
| At5g13330 | AP2/EREBP | RAP2.6L | 0.63 | 0.05 | **2.08** | 0.16 |
| At1g48150 | MADS | AgL74 | 0.76 | 0.09 | **1.93** | 0.17 |
| At2g22770 | bHLH | NAI | 0.56 | 0.48 | **1.81** | 0.75 |
| At3g23230 | AP2/EREBP | ERF98 | -0.16 | 0.08 | **1.78** | 0.76 |
| At2g38340 | AP2/EREBP | DREB19 | 0.31 | 0.13 | **1.75** | 0.13 |
| At5g23000 | MYB | MYB37 | 0.87 | 0.33 | **1.69** | 0.46 |
| At1g62975 | bHLH | | 0.12 | 0.45 | **1.62** | 0.40 |
| At1g25470 | AP2/EREBP | CRF12 | 0.41 | 0.89 | **1.58** | 0.14 |
| At2g27220 | HB | BLH5 | -0.47 | 0.14 | **-1.57** | 0.69 |
| At2g25820 | AP2/EREBP | ESE2 | -0.21 | 0.20 | **-1.65** | 0.22 |
| At3g58070 | C2H2 | glS | -0.46 | 0.41 | **-1.87** | 0.12 |
| At3g46130 | MYB | MYB48 | -0.23 | 0.19 | **-1.97** | 0.22 |
| At3g15320 | MYB | | -0.70 | 0.05 | **-2.00** | 0.64 |
| At3g53680 | PHD finger | | -0.41 | 0.04 | **-2.06** | 0.15 |
| At5g51870 | MADS | AgL71 | -0.68 | 0.14 | **-2.09** | 0.20 |
| At4g22950 | MADS | AgL19 | 0.58 | 0.63 | **-2.48** | 0.11 |

**Supplementary Table III. Expression of TF-encoding genes modulated by absence of both MKP1 and AP2C1.** Transcript levels were quantified by RT-qPCR and are expressed as fold change (FC) in $log_2$ scale for each of the mutants compared to WT. Genes affected by at least 1.56 both in *ap2c1* and *mkp1* are listed. Data are from 2 independent biological replicates and are reported with SE.

| AGI | gene family | gene name(s) | *ap2c1* $log_2$ FC | SE of -ΔΔct | *mkp1* $log_2$ FC | SE of -ΔΔct |
|---|---|---|---|---|---|---|
| At3g47870 | AS2 (LOB) I | ASL29/LBD27/SCP | **2.24** | 0.25 | **3.57** | 0.07 |
| At2g13150 | bZIP | BZIP31 | **2.25** | 0.77 | **3.76** | 0.76 |
| At1g03790 | C3H | SOM/TZF4 | **2.27** | 0.15 | **1.81** | 0.49 |
| At1g18860 | WRKY | WRKY61 | **1.99** | 0.11 | **2.49** | 0.08 |

**Supplementary Table IV. TF-encoding genes de-regulated in *ap2c1 mkp1* plants**

Gene transcript levels were quantified by RT-qPCR and expressed as fold change in $log_2$ scale for each of the mutants compared to WT. Genes affected in *ap2c1 mkp1* compared to WT by at least 1.56 in $log_2$ scale are listed. Data are from 3 independent biological replicates and are reported with SD.

**upregulated in *ap2c1 mkp1***

| | | *ap2c1* | | *mkp1* | | *ap2c1 mkp1* | |
|---|---|---|---|---|---|---|---|
| AGI | family | -ΔΔct | SD | -ΔΔct | SD | -ΔΔct | SD |
| At5g13330 | AP2/EREBP | 0,53 | 0,15 | 2,33 | 0,37 | 5,79 | 1,77 |
| At1g36060 | AP2/EREBP | -0,60 | 0,50 | 1,77 | 0,09 | 4,60 | 0,95 |
| At3g23240 | AP2/EREBP | 1,02 | 0,11 | 2,65 | 0,20 | 4,44 | 1,85 |
| At2g38340 | AP2/EREBP | 0,54 | 0,37 | 1,81 | 0,39 | 4,43 | 1,35 |
| At1g04370 | AP2/EREBP | 1,57 | 0,24 | 1,35 | 0,77 | 4,43 | 2,11 |
| At2g47520 | AP2/EREBP | 0,78 | 0,81 | 0,22 | 0,41 | 4,06 | 1,50 |
| At5g64750 | AP2/EREBP | 0,46 | 0,91 | 1,13 | 0,72 | 2,38 | 0,33 |
| At2g33710 | AP2/EREBP | 0,59 | 0,50 | 3,09 | 0,84 | 5,56 | 0,83 |
| At1g71520 | AP2/EREBP | 0,41 | 0,91 | 3,71 | 0,54 | 5,05 | 1,33 |
| At3g23230 | AP2/EREBP | -0,16 | 0,08 | 1,91 | 1,24 | 3,58 | 0,40 |
| At2g40470 | AS2 (LOB) I | 0,31 | 0,94 | 1,72 | 0,51 | 4,30 | 1,10 |
| At1g67100 | AS2 (LOB) II | 0,79 | 1,00 | 1,54 | 0,37 | 5,91 | 1,92 |
| At5g43650 | bHLH | -0,03 | 0,65 | 2,22 | 2,20 | 5,88 | 1,94 |
| At2g43140 | bHLH | 0,17 | 0,14 | 1,78 | 0,24 | 4,02 | 0,79 |
| At5g56960 | bHLH | -0,14 | 0,25 | 1,94 | 1,21 | 5,62 | 0,81 |
| At1g08320 | bZIP | 0,93 | 0,64 | 1,87 | 0,25 | 4,15 | 1,14 |
| At3g53600 | C2H2 | 0,59 | 1,13 | 1,38 | 1,84 | 4,58 | 2,56 |
| At5g67450 | C2H2 | -0,12 | 0,48 | 0,64 | 0,35 | 3,33 | 1,15 |
| At5g04390 | C2H2 | 0,15 | 0,62 | -0,22 | 0,93 | 2,88 | 0,82 |
| At2g37430 | C2H2 | 0,21 | 0,66 | 0,33 | 0,73 | 2,68 | 0,27 |
| At3g46080 | C2H2 | 0,21 | 0,65 | 0,80 | 0,57 | 7,15 | 1,26 |
| At3g46090 | C2H2 | -0,77 | 0,40 | 0,16 | 0,55 | 5,92 | 0,92 |
| At5g59820 | C2H2 | 0,54 | 1,14 | 0,50 | 0,84 | 4,14 | 0,85 |
| At1g13300 | GARP-G2-like | 0,38 | 1,09 | 0,46 | 1,17 | 4,50 | 0,96 |
| At5g19520 | HB | 0,91 | 0,91 | 0,02 | 1,56 | 2,47 | 0,38 |
| At4g36990 | HSF | 0,63 | 0,44 | 0,31 | 1,13 | 4,62 | 1,21 |
| At4g18870 | HSF | 3,27 | 0,83 | 1,24 | 1,10 | 7,52 | 2,84 |
| At5g20240 | MADS | 0,02 | 0,88 | 0,56 | 0,95 | 4,86 | 1,06 |
| At5g23000 | MYB | 0,87 | 0,33 | 1,77 | 0,40 | 4,26 | 1,44 |
| At1g18570 | MYB | 0,21 | 0,50 | 0,60 | 0,54 | 2,62 | 0,70 |
| At1g74080 | MYB | 0,21 | 0,58 | 0,61 | 1,06 | 8,00 | 1,78 |
| At3g23250 | MYB | 0,93 | 0,94 | 0,45 | 0,72 | 4,51 | 1,75 |
| At1g02250 | NAC | -0,68 | 0,15 | 0,23 | 0,51 | 5,57 | 1,69 |
| At1g02220 | NAC | 0,34 | 0,90 | 0,52 | 0,50 | 4,65 | 0,93 |
| At3g04070 | NAC | 0,84 | 0,62 | 1,57 | 0,67 | 4,32 | 0,86 |
| At3g15500 | NAC | 0,90 | 0,70 | 2,28 | 1,11 | 3,88 | 1,29 |
| At1g01010 | NAC | -0,03 | 0,38 | -0,02 | 0,62 | 3,12 | 0,63 |
| At3g44350 | NAC | -0,41 | 0,62 | -0,78 | 0,59 | 3,09 | 1,05 |
| At2g43000 | NAC | 0,78 | 0,62 | 3,16 | 0,83 | 4,94 | 1,89 |
| At2g38250 | Trihelix | -0,01 | 0,68 | 0,19 | 0,69 | 3,13 | 0,93 |
| At3g01970 | WRKY | 0,56 | 0,20 | 1,40 | 0,82 | 4,66 | 1,63 |
| At5g46350 | WRKY | 0,38 | 0,84 | 0,94 | 0,59 | 3,82 | 1,16 |
| At1g62300 | WRKY | 0,10 | 0,45 | 1,03 | 1,06 | 3,51 | 0,95 |
| At4g01720 | WRKY | 0,06 | 0,10 | 0,74 | 0,66 | 2,77 | 0,76 |
| At4g18170 | WRKY | 0,25 | 0,72 | 0,65 | 1,00 | 2,68 | 0,45 |
| At2g38470 | WRKY | -0,18 | 0,27 | 0,39 | 0,61 | 2,46 | 0,62 |
| At5g13080 | WRKY | 0,34 | 0,30 | 3,55 | 1,20 | 8,31 | 1,12 |
| At5g24110 | WRKY | 1,45 | 0,87 | 2,43 | 0,79 | 7,03 | 0,72 |
| At1g29860 | WRKY | 1,35 | 0,71 | 2,02 | 0,92 | 5,91 | 2,17 |
| At1g18860 | WRKY | 2,00 | 0,09 | 2,90 | 0,59 | 5,46 | 2,48 |
| At1g66560 | WRKY | 0,49 | 1,24 | 1,22 | 0,80 | 5,46 | 1,01 |
| At5g01900 | WRKY | 1,37 | 0,32 | 0,06 | 1,40 | 4,93 | 1,64 |
| At5g22570 | WRKY | 0,67 | 0,30 | 1,40 | 0,52 | 4,92 | 1,54 |
| At5g64810 | WRKY | 0,22 | 0,50 | -0,07 | 1,37 | 4,74 | 1,77 |
| At4g23810 | WRKY | 0,40 | 0,20 | 0,32 | 0,76 | 4,54 | 0,97 |
| At2g40740 | WRKY | 1,22 | 0,59 | 0,07 | 0,81 | 4,13 | 1,54 |
| At1g80840 | WRKY | 0,45 | 1,04 | 1,09 | 0,91 | 3,66 | 0,85 |
| At1g69600 | ZF-HD | 0,58 | 0,82 | -0,18 | 0,93 | 4,12 | 1,76 |

**downregulated in *ap2c1 mkp1***

| | | *ap2c1* | | *mkp1* | | *ap2c1 mkp1* | |
|---|---|---|---|---|---|---|---|
| AGI | family | -ΔΔct | SD | -ΔΔct | SD | -ΔΔct | SD |
| At3g26790 | ABI3/VP1 | -0,49 | 0,06 | -5,42 | 1,71 | -6,84 | 2,39 |
| At3g50510 | AS2 (LOB) I | -0,09 | 1,26 | -8,09 | 0,51 | -2,15 | 0,34 |
| At4g32280 | Aux/IAA | 0,60 | 1,17 | -0,88 | 0,89 | -3,57 | 0,47 |
| At1g52830 | Aux/IAA | 0,11 | 0,46 | -1,46 | 0,36 | -2,89 | 0,58 |
| At1g04240 | Aux/IAA | -0,16 | 0,58 | -0,83 | 0,48 | -2,48 | 0,65 |
| At2g01200 | Aux/IAA | -0,04 | 0,48 | -0,40 | 0,91 | -1,83 | 0,67 |
| At3g21330 | bHLH | 0,20 | 0,43 | -1,32 | 1,22 | -4,06 | 0,83 |
| At1g02340 | bHLH | 0,75 | 0,82 | 0,18 | 1,00 | -3,31 | 1,17 |
| At5g51790 | bHLH | 0,46 | 0,41 | 0,14 | 0,25 | -2,69 | 1,13 |
| At5g39860 | bHLH | 0,56 | 0,82 | -0,22 | 0,64 | -2,06 | 0,15 |
| At5g15160 | bHLH | -0,09 | 0,19 | -0,61 | 0,41 | -2,03 | 0,68 |
| At5g44260 | C3H | 0,67 | 0,94 | -0,71 | 0,98 | -2,93 | 0,69 |
| At3g13840 | GRAS | -0,96 | 0,19 | -0,12 | 0,50 | -2,12 | 0,72 |
| At5g27810 | MADS | -0,07 | 1,11 | -2,10 | 0,56 | -3,32 | 1,03 |
| At3g15320 | MYB | -0,35 | 0,50 | -2,39 | 0,75 | -3,89 | 1,03 |
| At5g56840 | MYB-related | 0,01 | 0,73 | -0,53 | 1,35 | -3,61 | 1,77 |
| At1g75250 | MYB-related | -0,31 | 0,53 | -1,09 | 0,20 | -2,47 | 0,81 |
| At5g43290 | WRKY | 0,97 | 0,00 | -1,32 | 0,40 | -3,89 | 1,68 |

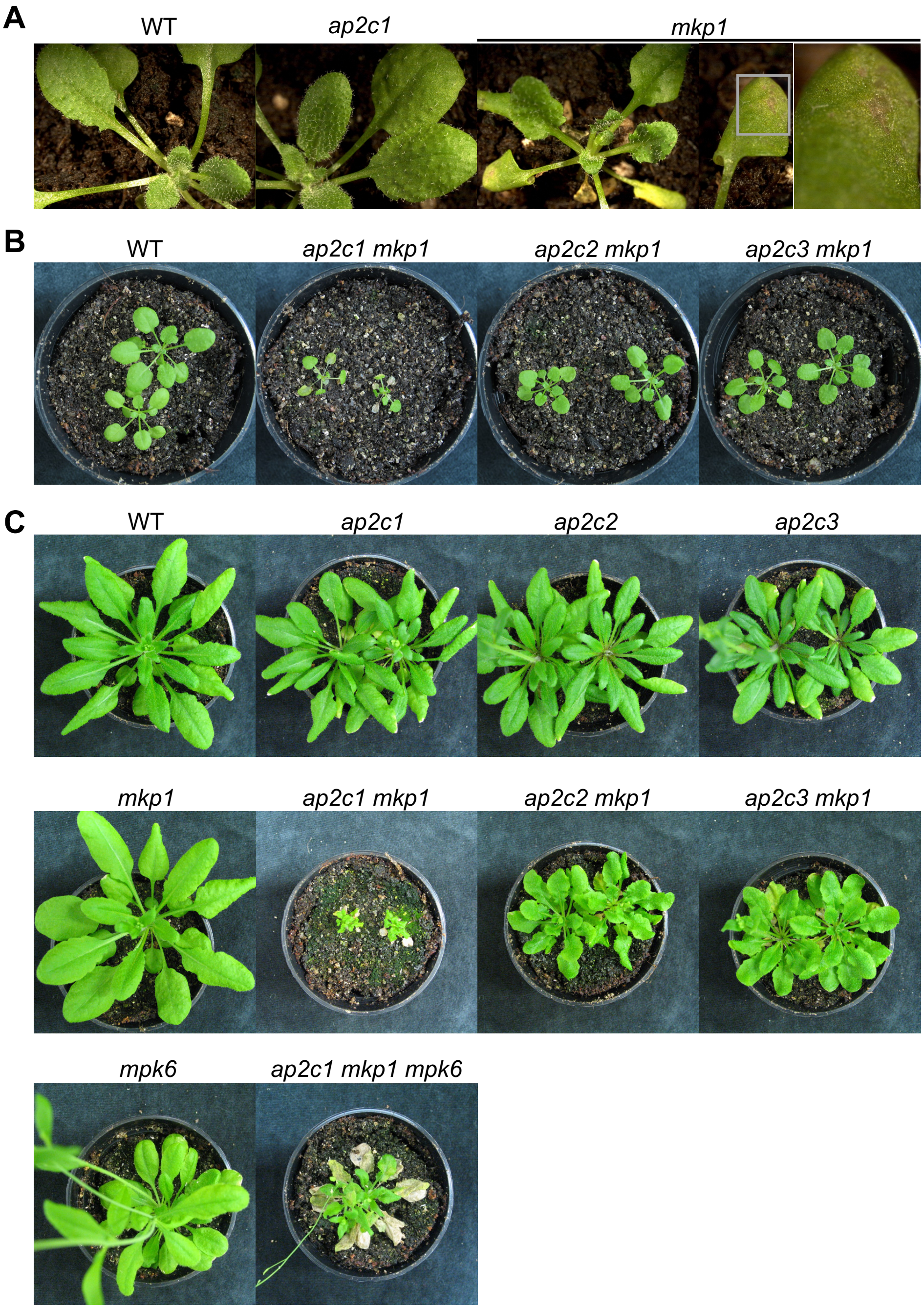


**Supplementary Figure 1. Phenotypes of Arabidopsis single, double and triple mutant plants.**

**A.** Phenotypes of WT, *ap2c1,* and *mkp1* plants grown for three weeks in long day conditions.

**B.** Phenotypes of WT, *ap2c1 mkp1, ap2c2 mkp1,* and *ap2c3 mkp1* plants grown for four weeks in short-day conditions.

**C.** Phenotypes of WT, *ap2c1, ap2c2, ap2c3, mkp1, ap2c1 mkp1, ap2c2 mkp1, ap2c3 mkp1, mpk6,* and *ap2c1 mkp1 mpk6* plants grown for four weeks in short-day condition, followed by three weeks in long-day condition.

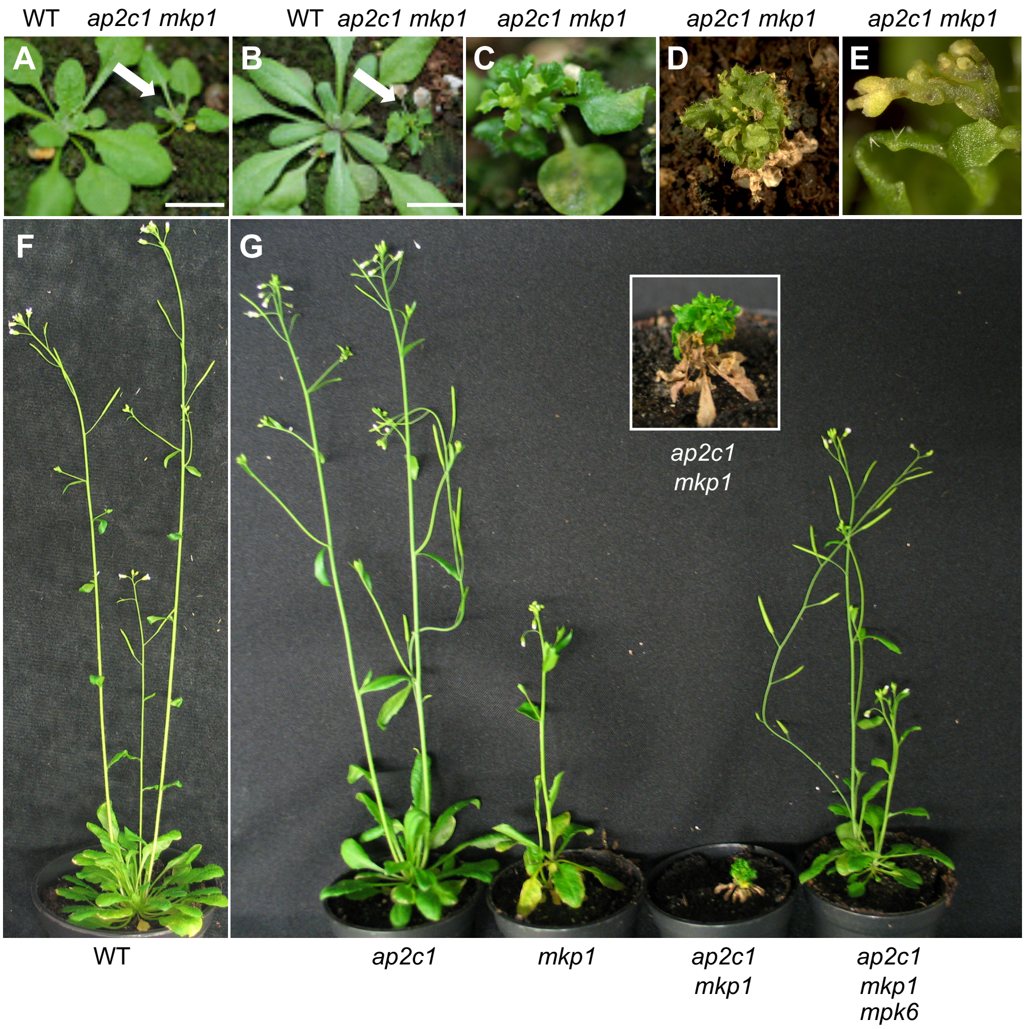


**Supplementary Figure 2. Loss of both AP2C1 and MKP1 leads to severe phenotypes in growth and development, which are mediated by MPK6.**

**A.** Phenotypes of WT (left) and *ap2c1 mkp1* (right, indicated by arrow) grown 2.5 weeks in standard long-day conditions. During the first 18 days of growth the phenotypic differences of *ap2c1 mkp1* compared to WT plants are visibly manifested as a difference in plant size.

**B.** After approximately 3.5 weeks (26 days) premature death of leaf tissue as well as abnormal leaf growth and morphology of *ap2c1 mkp1* (right, indicated by arrow) become apparent. Scale bars in **A** and **B** = 1 cm.

**C.** Close-up of *ap2c1 mkp1* plant 3.5 weeks grown in standard long-day conditions.

**D.** Phenotype of seven-week-old *ap2c1 mkp1* plant grown in standard long-day condition.

**E.** Close-up of seven-week-old *ap2c1 mkp1* plant showing misshaped inflorescence.

**F, G**. Phenotypes of eight-week-old WT, *ap2c1*, *mkp1*, *ap2c1 mkp1* and *ap2c1 mkp1 mpk6* plants grown the first six weeks in short-day and for a further two weeks in long-day conditions. The *ap2c1 mkp1* double mutant display a severe dwarf phenotype, premature leaf decay, lack of normal shoot development, and strongly impaired inflorescence growth. The inset picture above shows a close-up of *ap2c1 mkp1* plant shown below. In *ap2c1 mkp1 mpk6* triple mutant plants these phenotypes were rescued.
